# BionetBF: A Novel Bloom Filter for Faster Membership Identification of Large Biological Graph

**DOI:** 10.1101/2021.09.23.461527

**Authors:** Sabuzima Nayak, Ripon Patgiri

**Affiliations:** Department of Computer Science & Engineering, National Institute of Technology Silchar, Silchar, 788010, Assam, India

**Keywords:** Bloom Filter, Big Graph, Biological Network, Probabilistic Data Structure, Algorithm

## Abstract

Big Graph is a graph having thousands of vertices and hundreds of thousands of edges. The study of graphs is crucial because the interlinkage among the vertices provides various insights and uncovers the hidden truth developed due to their relationship. The graph processing has non-linear time complexity. The overwhelming number of vertices and edges of Big Graph further enhances the processing complexity by many folds. One of the significant challenges is searching for an edge in Big Graph. This article proposes a novel Bloom Filter to determine the existence of a relationship in Big Graph, specifically biological networks. In this article, we propose a novel Bloom Filter called Biological network Bloom Filter (BionetBF) for fast membership identification of the biological network edges or paired biological data. BionetBF is capable of executing millions of operations within a second while occupying a tiny main memory footprint. We have conducted rigorous experiments to prove the performance of BionetBF with large datasets. The experiment is performed using 12 synthetic datasets and three biological network datasets. It takes less than 8 sec for insertion and query of 40 million biological edges. It demonstrates higher performance while maintaining a 0.001 false positive probability. BionetBF is compared with other filters: Cuckoo Filter and Libbloom, where small-sized BionetBF proves its supremacy by exhibiting higher performance compared with large-sized Cuckoo Filter and Libbloom. The source code is available at https://github.com/patgiri/BionetBF. The code is written in the C programming language. All data are available at the given link.

**Highlights:**

- Proposed a novel Bloom Filter, BionetBF, for faster boolean query on Big Graph.
- BionetBF has a low memory footprint and the lowest false positive probability.
- It has high performance with constant searching time complexity.
- BionetBF has the potential to application in Big Graph, de-Bruijn Graph, and Drug Discovery.

## 1. Introduction

The graph has always been the most crucial research area for years due to its diverse application fields. The graph helps in the representation of relationships among the entities. For example, social networks represent the connectivity of users based on friendship. The second attractive feature of graphs is non-linear time complexity. Every operation in a graph, for instance, insertion, query and deletion, has non-linear time complexity. It excites the researcher to develop an algorithm that performs the operations in linear time. Today’s digital era has converted the graph into Big Graph. The exponential generation of data has led to Big Graphs. For example, the number of social network users is millions, and each user is connected to thousands of other users. The social network graph has millions of vertices with a hundred million edges. The number of monthly active users on Facebook is 2.98 billion in the year 2022 [1]. Big Graph has vast applications. Social network analysis helps in sociology. Sociology aims to understand society, patterns of social relationships, human social behaviour, social interaction, etc. Knowledge graph [2, 3] presents the data and their relationship, for example, google knowledge graph [4]. Big Graph is big, complex, uncertain, and dynamic [5]. These features influence the design of a Big Graph searching algorithm. The colossal number of vertices and edges increases the complexity and uncertainty in the Big Graph. It increases the processing complexity by many folds. Due to the huge size, the searching algorithm has to balance the time and space complexity. The algorithm design should be a reasonable model to capture the uncertainties of the Big Graph. Many Big Graphs are dynamic. New edges or vertices are added or deleted from the graph. For example, a user frequently adds friends on social networks and changes the graph structure. The searching algorithm needs to be fast to provide a result before the input graph structure becomes invalid. Another challenge associated with the Big Graph is the inability to store the entire graph in RAM. One solution is distributed graph processing, for example, Pregel [6]. Pregel has scalability issues such as superstep barriers [7] and network traffic issues in case of dense graphs. Bloom Filter [8] is a probabilistic data structure which is not a complete system but can reduce the burden of the state-of-the-art techniques in graph searching to a large extent.

A biological network is a complex network that represents the relationship between biological entities such as molecules, proteins, ions, metabolites, etc. [9]. Some examples of biological network are protein-protein interaction networks [10], genetic regulatory networks, cell-cell communication [11], and genetic interaction networks [12]. Protein-protein interaction networks present transient and stable interactions between the proteins where the interactions are undirected by nature [13]. The gene regulatory network represents the complete set of gene products and their interactions. Genetic interaction networks represent the interactions between the genes within an organism [14, 15]. Moreover, Bloom Filter is used to construct de Bruijn Graph where de Bruijn Graph of human genome size is more than 30GB of RAM [16, 17]. Therefore, Bloom Filter is an efficient option to use to solve the de-Bruijn graph problem.

Representing the biological system as a network instance increases the comprehension of the complex biological process, particularly at the molecular level [18]. Biological networks unfold prominent relationships, patterns, and properties of the whole system, which is difficult in a univariate analysis [19]. The study on the biological network of proteinprotein interaction helps determine the protein related to the diseases [20, 21]. The interactions among the proteins result in cellular and molecular mechanisms determining an organism’s health. A biological network is a great help in drug discovery. Typically, the commercial availability of a drug takes more than 7 years, the development of a drug takes approximately 5 years, and a few years more for clinical trials. Thus, the process of drug discovery is expensive and time-consuming. A biological network helps in discovering the valuable information related to a drug and its target, such as (a) determining any secondary effects interfering with the target, (b) whether the target protein has many connections, (c) whether the proteins of the targeted drug is dangerous, and (d) determines the molecules affected by the side effects of the drug. Furthermore, a biological network is used for predicting the drug combination because the network is fitting for computational and combinatorial analysis. The study of genetic interaction networks helps comprehend the relationship between phenotype and genotype. Phenotype is an organism’s trait resulting from the interaction of the genotype and its environment. The genotype is the collection of genes. The biological network also helps in determining how viruses and bacteria infect, persevere, and cause diseases [22, 23]. In addition, it helps to apprehend the pharmacokinetic and pharmacogenomic actions of antibacterial drugs. Furthermore, a biological network helps establish the relationships among organisms and species [24].

### 1.1. Motivation

Analysis of large networks is predominantly computationally intractable [25]. In other words, computationally intractable problems are either NP-hard or NP-complete problems that are solved approximately or heuristically. The various factors contributing to the complexity of biological networks are chemical kinetics, a vast number of interacting variables, feedback loops, dynamic in nature caused due to various linear or nonlinear relationships, and stochasticity [26]. The biological network is dynamic in nature [27]; however, the information provided by the biological network is static at an instance. The biological network is highly influenced by external factors making it dynamic in nature. Comparatively, a biological network furnishes a small part of the whole biological network; hence, it is missing much information. The biological network has thousands of millions of vertices with billions of interactions/edges. The huge amount of information the biological network generates often yields dubious interaction. The construction of a biological network is challenging due to the large biological data silo, and the data are noisy [28]. Thus, it demands highly efficient data structures with low memory footprints.

### 1.2. Contribution

This article proposes a novel Bloom Filter, called BionetBF, for rapid membership identification of edges in Big Graph. To the best of our knowledge, no Bloom Filter or filters are proposed for storing the Big Graph in a single data structure. We have considered biological networks for our experimentation. BionetBF can perform insertion and query operations on millions of biological edges within a few seconds. We have conducted extensive experiments on BionetBF using 12 synthetic biological datasets and 3 real datasets. In the case of synthetic biological datasets, BionetBF takes 7.6 sec for the insertion of 40 million biological edges. Similarly, it takes 7.6 sec for the query of 40 million biological edges. BionetBF maintains a low false positive probability of 0.001. One important point to notice is that BionetBF has zero false positive probability in many datasets. The performance of BionetBF is measured using multiple parameters: Million operations per second (MOPS), Seconds per operation (SPO), and Megabyte per operation (MBPS). In insertion operation, the highest MOPS, SPO, and MBPS are 7.06, 1.87, and 162.78, respectively. Similarly, in query operation (Disjoint set), the highest MOPS, SPO, and MBPS are 6.95, 1.9, and 160.6, respectively. BionetBF demonstrates an accuracy of more than 99.5%. BionetBF is compared with another two filters: Cuckoo Filter [29] and Libbloom (standard Bloom Filter) [30]. A small-sized BionetBF demonstrates faster operations and high performance than Cuckoo Filter and Libbloom. Cuckoo Filter requires 6.6×, and Libbloom requires 4.7× larger memory than BionetBF. Furthermore, the performance of BionetBF is measured using three biological network datasets: Drug-Gene, Gene-Disease, and Gene-Gene. Drug-Gene is the dataset of interaction between molecules and genes. Gene-Disease is the dataset about the association of disease with genes. Gene-Gene is the dataset about the interaction among genes. BionetBF also presents a higher performance in the biological network datasets. BionetBF can be used for quick identification of a specific biological edge in many biological networks. BionetBF is also an excellent choice for filtering repetitive insertion or query of biological edge.

## 2. Bloom Filter

### Definition 1.

*Bloom Filter is an approximation data structure to test whether a queried item is a member of a set or not. It returns either definitely not in the set or possibly in the set. An empty Bloom filter is a bit array of m bits, all set to 0.It requires k independent hash functions to map an input item into Bloom Filter. In insertion, all corresponding k bit positions of the bit array are set to 1 using k hash functions. In a query operation, all k bit positions of the bit array must be 1 to return true; otherwise, Bloom Filter returns false, which refers that the queried item is not in the set [31]*.

Bloom Filter [8] is an approximate set membership filtering data structure defined in Definition 1. It is extremely popular in diverse domains, particularly, Big Data [32], IoT, Cloud Computing, Networking [33], Security [34, 35], Database, Bioinformatics [36], and Biometrics. Bloom Filter is applied to reduce the main memory footprint. Bloom Filter uses a tiny amount of main memory to filter mammoth-sized data. It can significantly boost the performance of any system, but it has a false positive probability issue. Nevertheless, designing a new Bloom Filter can mitigate the false positive probability.

Standard Bloom Filter/Bloom Filter is a bit array of *m* bits. It uses *k* number of hash functions. Initially, each bit/cell is set to 0. Bloom Filter performs two operations: insertion and query. In insertion operation, *k* hash functions hash a single item. The hash value gives a location of a cell. The *k* cells are set to 1. Similarly, in query operation, the *k* hash functions hash the queried item. If all the *k* cells are 1, then Bloom Filter returns true. If at least one cell is 0, then Bloom Filter returns false.

Bloom Filter [8] is a bitmap array consisting of bits. Let 𝔹 be a Bloom Filter of size *m*, 𝕊 = {*e*_1_, *e*_2_, *e*_3_, …, *e*_*n*_} be the set of words inserted into the Bloom Filter 𝔹, U be the universe where 𝕊 ⊂ U, *k* is the number of hash functions. Bloom Filter size (i.e., *m*) is a prime number to reduce collision [31]. Bloom Filter calculates the optimal number of hash functions using Equation 1.

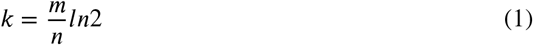

A large *k* value increases the time complexity of the operations, whereas a small *k* value increases collision probability. Bloom Filter returns either true or false in query operation, but these responses are further classified based on the presence/absence of the words in Bloom Filter. The *True* is classified into *True Positiυe* and *True Negatiυe*, and *False* is classified into *False Positiυe* and *False Negatiυe*. Let *e*_*i*_ be the random query item and queried to the Bloom Filter, then the definitions are as follows-

### Definition 2.

*If e*_*i*_ ∈ 𝕊, *and e*_*i*_ ∈ 𝔹, *then the result of* 𝔹 *is true positive*.

### Definition 3.

*If e*_*i*_ ∉ 𝕊, *but e*_*i*_ ∈ 𝔹, *then the result of* 𝔹 *is false positive*.

### Definition 4.

*If e*_*i*_ ∉ 𝕊, *and e*_*i*_ ∉ 𝔹, *then the result of* 𝔹 *is true negative*.

### Definition 5.

*If e*_*i*_ ∈ 𝕊, *but e*_*i*_ ∉ 𝔹, *then the result of* 𝔹 *is false negative*.

Figure 1 illustrates the bit array of a standard Bloom Filter. The Bloom Filter in the Figure 1 has *k* = 3, i.e., *h*_1_(), *h*_2_(), and *h*_3_(), and X, Y, Z and Q are words. Initially, the bit array of Bloom Filter is set to 0. The word X is inserted into the Bloom Filter and is hashed by three hash functions. The hash value gives the bit location as pointed in Figure 1. Three hash functions give three bit locations which are set to 1. Follows a similar procedure to insert Y. In query operation, the same hash functions used in insertion operation hash the query word to obtain the bit locations. Checks the bit locations, whether all locations have the bit value as 1. If all bit locations are 1, the Bloom Filter returns *True*; if at least one bit location is 0, then Bloom Filter returns *False*. The query of X gives *True* response, whereas the query of Z gives a False response. In the case of the query of Z, out of three bit locations, one bit location is 0; hence, Bloom Filter returns *False*.

**Figure 1:**
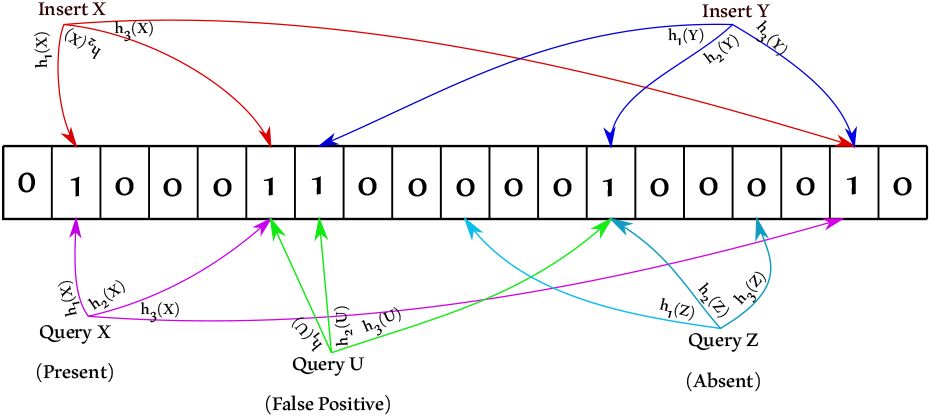
Architecture of Standard Bloom Filter using three hash functions.

### Algorithm 1

Insertion of a word *Word* into Bloom Filter 𝔹 using three hash functions as an example. The seed values *Seed S*_1_, *S*_2_, *S*_3_ are used to create different hash functions.

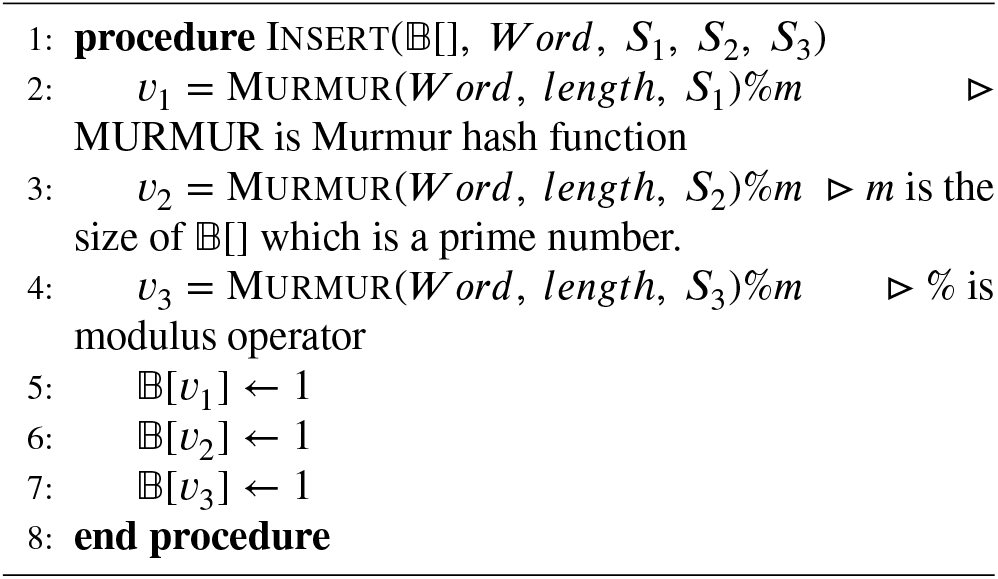

Bloom Filter has mainly two operations: insertion and uery. Algorithm 1 and 2 represent the insertion, and query operations, respectively. For example, the algorithms are written using the murmur hash function [37] with *k* = 3. Algorithm 1 inserts a word *Word* into Bloom Filter 𝔹. The modulus operation is performed on the hash value to limit the value within the size of the 𝔹. Modulus operation gives three bit locations which are set to 1. Algorithm 2 queries a word *Word* into 𝔹. It follows a procedure similar to the insertion operation to obtain the bit locations. Then, it performs *AND* operations to determine whether all bit locations are 1. If all bit locations are 1, then return *True*; otherwise *False*.

### Algorithm 2

Query a word *Word* in Bloom Filter 𝔹[] using three hash functions as an example. The seed values *Seed S*_1_, *S*_2_, *S*_3_ are used to create different hash functions.

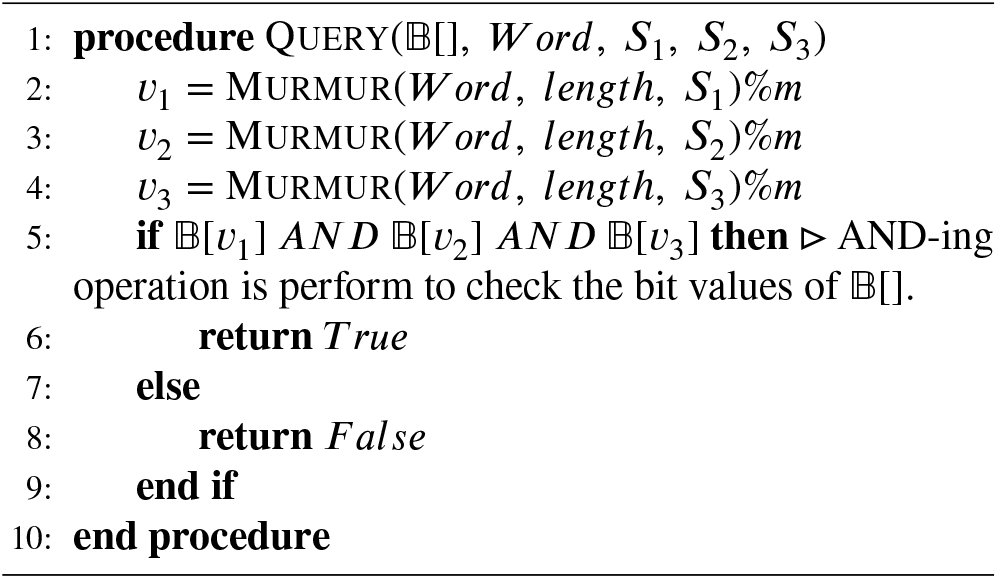

There is a prominent issue in Bloom Filter, particularly false positives. Assuming only words *X* and *Y* are inserted into 𝔹. A word *U* is queried to 𝔹 (as illustrated in Figure 1). 𝔹 hashes *U* by three hash functions. The three hash functions obtained three bit locations. All three locations have bit value Hence, 𝔹 returns *True*; which is an incorrect response. This is caused because the locations obtained are already set to 1 by the words *X* and *Y*.

## 3. Related Work

There are many variants of Bloom Filter such as Counting Bloom Filter [38], rDBF [39], Stateful Bloom Filter [40], etc. Moreover, many filters are proposed, for example, Tinyset [41], Xor Filter [42], Cuckoo Filter [29], etc.

Cuckoo Filter [29] is a data structure used for membership inspection, and it is not a Bloom Filter; instead, it is a filter performing the same task as Bloom Filter, i.e., membership identification. It uses a modified cuckoo hashing [43], called partial-key Cuckoo hashing. This hashing improves the dynamic insertion of words into the Cuckoo Filter. Instead of hash values, the fingerprint of the word is stored in the Cuckoo Filter to reduce memory requirements. The fingerprint size is less compared to the hashed value of the word. Cuckoo Filter generates two locations by hashing the fingerprint. Randomly any one location is selected for the storage of the fingerprint. In case the location is occupied, then select the other location. If both locations have fingerprints, then among the two locations, randomly select any one location. Then kick the fingerprint in the selected location and store the new fingerprint. Again, check the kicked fingerprint’s alternate location; if empty, store the kicked fingerprint; otherwise, repeat the kicking process. The kicking process continues until a threshold value. Crossing the kicking threshold confirms that the Cuckoo Filter is saturated. In query operation, hash the queried word to obtain the fingerprint. Again hash the fingerprint to generate two locations; if it is present in any one location, then the Cuckoo Filter returns True; otherwise, False.

## 4. Method

### 4.1. BionetBF: The Proposed System

BionetBF is a novel Bloom Filter for membership identification of edges in Big Graph. It is a two-dimensional Bloom Filter (2DBF) with fewer arithmetic operations. BionetBF is a two-dimensional array where the dimensions are different prime numbers. Prime numbers reduce collision probability. Usually, Bloom Filters take a single word as input, but the edge has two data/words (two vertices). Hence, the vertices of an edge are concatenated and given as input to BionetBF. This conversion of two words into a single word helps maintain a single data structure for the network dataset.

BionetBF implements two operations: insertion and query. Let a directed edge in a Big Graph be (*V*_1_, *V*_2_) where the edge direction is from *V*_1_ to *V*_2_. Our desired FPP is 0.001. BionetBF with one hash function has more than 0.001 FPP. Hence, BionetBF uses two hash functions, it is experimentally illustrated in the supplementary document. Let, 𝔹_*x,y*_ is a BionetBF, (*V*_1_, *V*_2_) is a directed edge, and ℋ() is a hash function. BionetBF uses the murmur hash function [37]. Different seed values in the murmur hash function create a different hash value for the same word. Initially, set all cells of BionetBF to 0.

Algorithm 3 demonstrates the insertion operation of BionetBF. First *V*_1_ and *V*_2_ are concatenated (say the concatenated word as *V*_1_*V*_2_ and *V*_1_*V*_2_ ≠ *V*_2_*V*_1_). The *V*_1_*V*_2_ is given as input to the murmur hash function. Then, it performs modular operations on the hashed value and the dimensions of the BionetBF. The result of the modular operations generates a location in BionetBF which stores the information indicating the insertion of the *V*_1_*V*_2_ into BionetBF. It performs *OR* operation to set one bit to 1 without losing previous information of that cell. This procedure is repeated twice as *k* = 2; in other words, each *V*_1_*V*_2_ sets two bits to 1 in BionetBF. In the case of an undirected edge, instead of concatenation *XOR* operation is performed between *V*_1_ and *V*_2_ and inserted into BionetBF. However, there is very less application of undirected edge. Hence, it is not explored in this article.

Algorithm 4 demonstrates the query operation of BionetBF for the queried item (*V*_1_, *V*_2_). The steps till obtaining a location in BionetBF are the same for both insertion and query operation. The XOR operation retrieves the related information from the cell, i.e., it gives the bit value of the queried edge. The *AND* operation determines whether the location bit is 1. It obtains two locations because *k* = 2. If both location bits are 1, then BionetBF returns True; otherwise, False.

#### Algorithm 3

Insertion of an edge (*V*_1_, *V*_2_) into BionetBF.

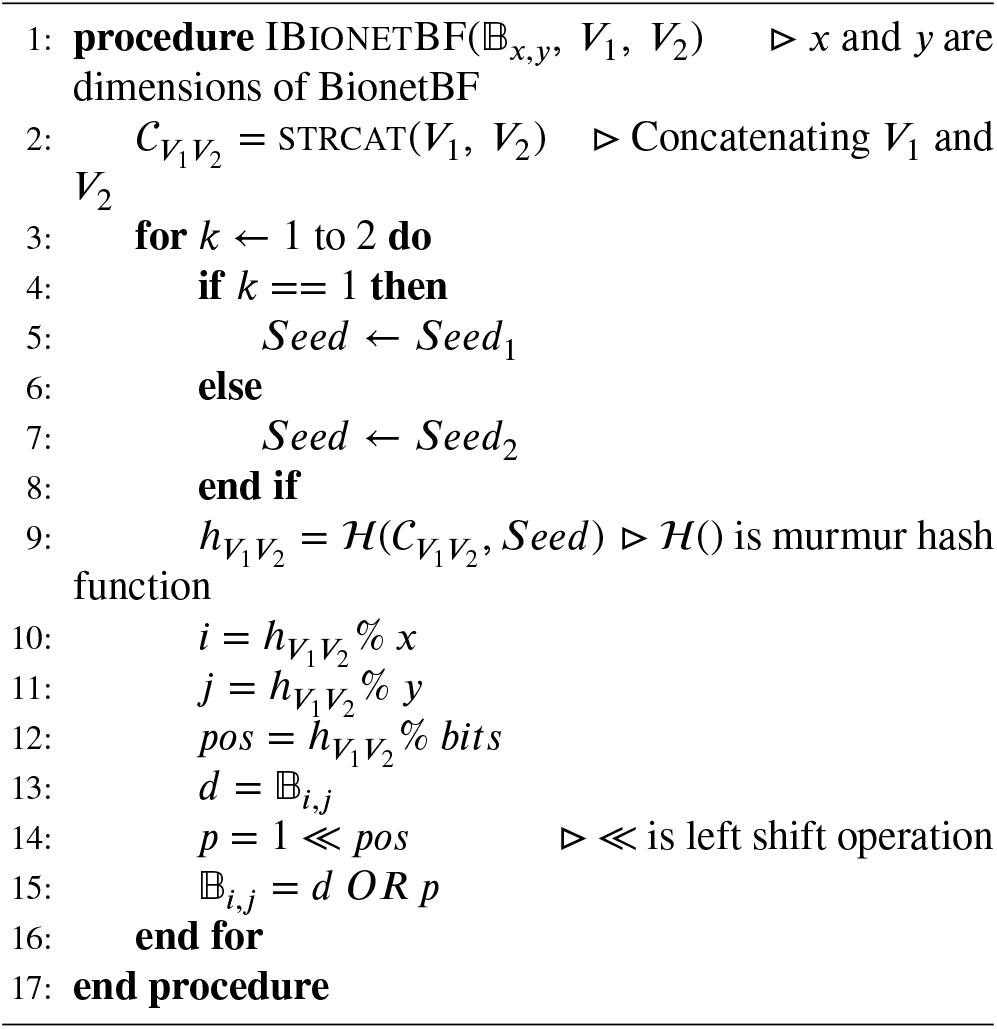

#### Algorithm 4

Query of an edge (*V*_1_, *V*_2_) into BionetBF.

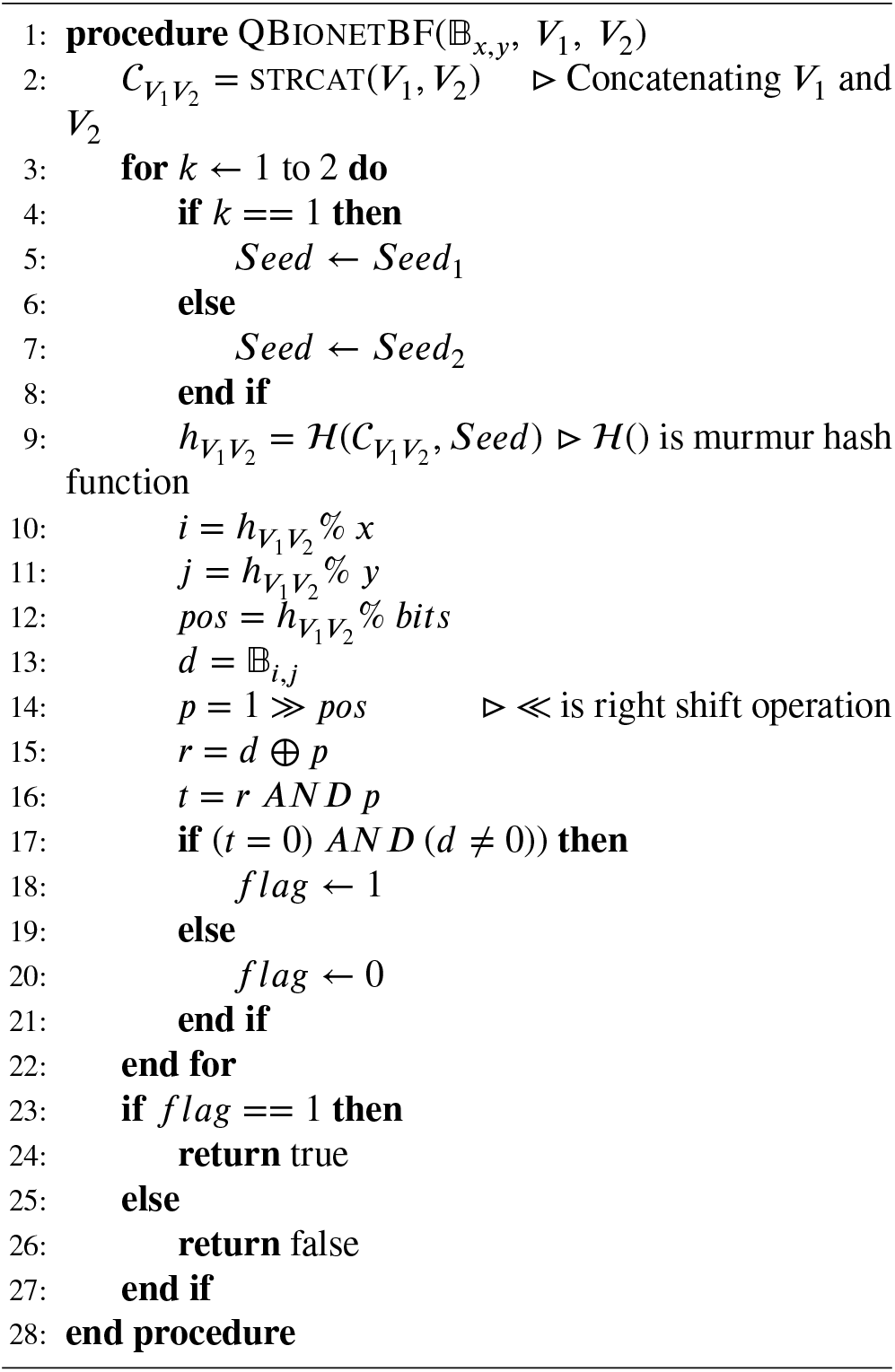

The presence of a path can also be verified in a Big graph by using performing multiple query operations in BionetBF. The *AND* operation is performed among individual query of all edges in the path. Lets demonstrates the path query operation with an example. Let (*V*_1_, *V*_2_, *V*_3_, *V*_4_) are four vertices that has a path from *V*_1_ to *V*_4_ such that *V*_1_ → *V*_2_, *V*_2_ → *V*_3_, and *V*_3_ → *V*_4_. Let us assume a query that whether their is a path from *V*_1_ to *V*_4_. The path can be queried into the Bloom Filter as 𝒫 = QBionetBF(𝔹_*x,y*_, *V*_1_, *V*_2_) *AND* QBionetBF(𝔹_*x,y*_, *V*_2_, *V*_3_) *AND* QBionetBF(𝔹_*x,y*_, *V*_3_, *V*_4_). If 𝒫 = *True*, then there exists a path, otherwise, not.

### 4.2. Data Description

We have used two types of datasets for the BionetBF experiments: synthetic and real-world datasets ^1^. The synthetic dataset is *IDKmer*, and the real-world dataset is the biological network dataset. We have used the synthetic dataset to verify the correctness of the experimental results. The real-world dataset is used to present the efficiency of BionetBF in real-world data. This section provides details of the datasets. To determine the FPP, we have generated three different datasets for query operation: Same Set 𝒮, Mixed Set ℳ and Disjoint Set 𝒟. Let the inserted data set in the Bloom Filter be ℐ (Inserted Set). The Same Set and Disjoint Set can be defined as ℐ = 𝒮 and 𝒟 ∩ 𝒮 = *ϕ*, respectively. Let, ℳ = {𝒬_1_, 𝒬_2_}, then one of the following condition is true for Mixed Set:

1. If 𝒬_1_ ∈ ℐ, then 𝒬_2_ ∈ 𝒟.
2. If 𝒬_1_ ∈ 𝒟, then 𝒬_2_ ∈ ℐ.

An example to illustrate the above definitions, Let ℐ = {(*υ*_1_, *υ*_2_), (*υ*_2_, *υ*_3_), (*υ*_3_, *υ*_4_), …, (*υ*_10_, *υ*_11_)} Then, 𝒮 = {(*υ*_1_, *υ*_2_), (*υ*_2_, *υ*_3_), (*υ*_3_, *υ*_4_), …, (*υ*_10_, *υ*_11_)} 𝒟 = {(*υ*_*a*_, *υ*_*b*_), (*υ*_*b*_, *υ*_*c*_), (*υ*_*c*_, *υ*_*d*_), …, (*υ*_*j*_, *υ*_*k*_)} and ℳ = {(*υ*_1_, *υ*_2_), (*υ*_2_, *υ*_3_), (*υ*_3_, *υ*_4_), (*υ*_4_, *υ*_5_), (*υ*_5_, *υ*_6_), (*υ*_*a*_, *υ*_*b*_), (*υ*_*b*_, *υ*_*c*_), (*υ*_*c*_, *υ*_*d*_), (*υ*_*d*_, *υ*_*e*_), (*υ*_*e*_, *υ*_*f*_)} Note: Only consider the set elements, not the structure of the graph.

The Same Set is not generated separately; instead, the ℐ is again queried to BionetBF. To avoid confusion between insertion and query set, the Same Set reference is used to indicate the queried set. Disjoint Set is a set having no common edge with the Inserted Set as depicted in Figure 2. The Mixed Set comprises half the number of edges of Inserted Set and Disjoint Set. The Same Set is used to determine the correct execution of the operation. Whereas Mixed Set and Disjoint Set are used to determine FPP in BionetBF.

**Figure 2:**
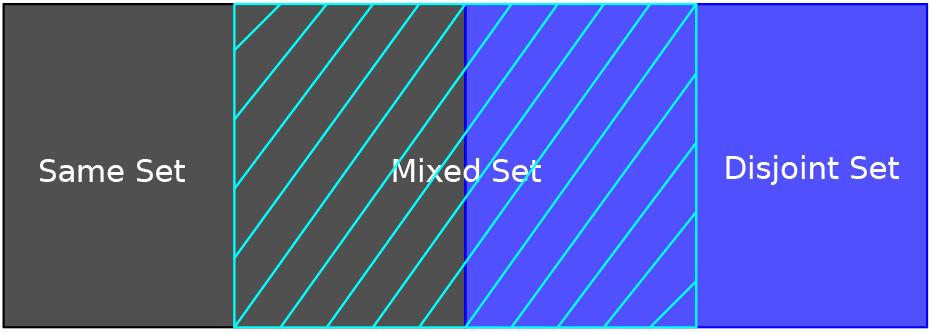
Dataset demonstration of Same Set, Mixed Set and Disjoint Set.

#### 4.2.1. IDKmer Dataset

The K-mer is defined as a DNA sequence or string of length *K*. Our synthetic dataset, *IDKmer*, consists of biological edge: *Kmer ID* and *K*−*mer*. The *K*−*mer* is a K-mer of a fixed length, and *Kmer ID* is the ID of the *K* −*mer*. For *K* − *mer*, read a genomic sequence of a fixed length from a DNA sequence dataset (downloaded from [dataset] consists of human DNA sequences [44]). The *K* − *mer* lengths considered for the experiment are 8, 15 and 20. The *K* −*mer* are read from the single sequence continuously, i.e., the reading of the first sequence starts from the first nucleotide of the DNA sequence dataset. The sequence is read for the required length. Then, for the second sequence, the reading of the nucleotide starts from the nucleotide, where it was stopped in the first sequence. To assign unique *Kmer ID*, an 8 digit long number is incremented and assigned to each sequence. Disjoint set is generated by taking a different 8 digit long number from the original set. There are four datasets with 10, 20, 30 and 40 million numbers of biological edge having *K*−*mer* length 8. Similarly, the IDKmer dataset with *K*−*mer* length 15 and 20 is generated; each has four datasets with 10, 20, 30 and 40 million biological edges. Hence, there are 12 *IDKmer* datasets. Each 12 *IDKmer* dataset has its Mixed and Disjoint sets. Table 1 lists the notation used to refer to the datasets and the file size. In our synthetic dataset, the first *IDKmer* biological edge with *K* − *mer* length 8 is “10000000 CTGGGCTA”, with Kmer length 15 is “10000000 CTGGGCTAAAAGGTC” and with K-mer length 20 is “10000000 CTGGGCTAAAAGGTCCCTTA”.

**Table 1.**
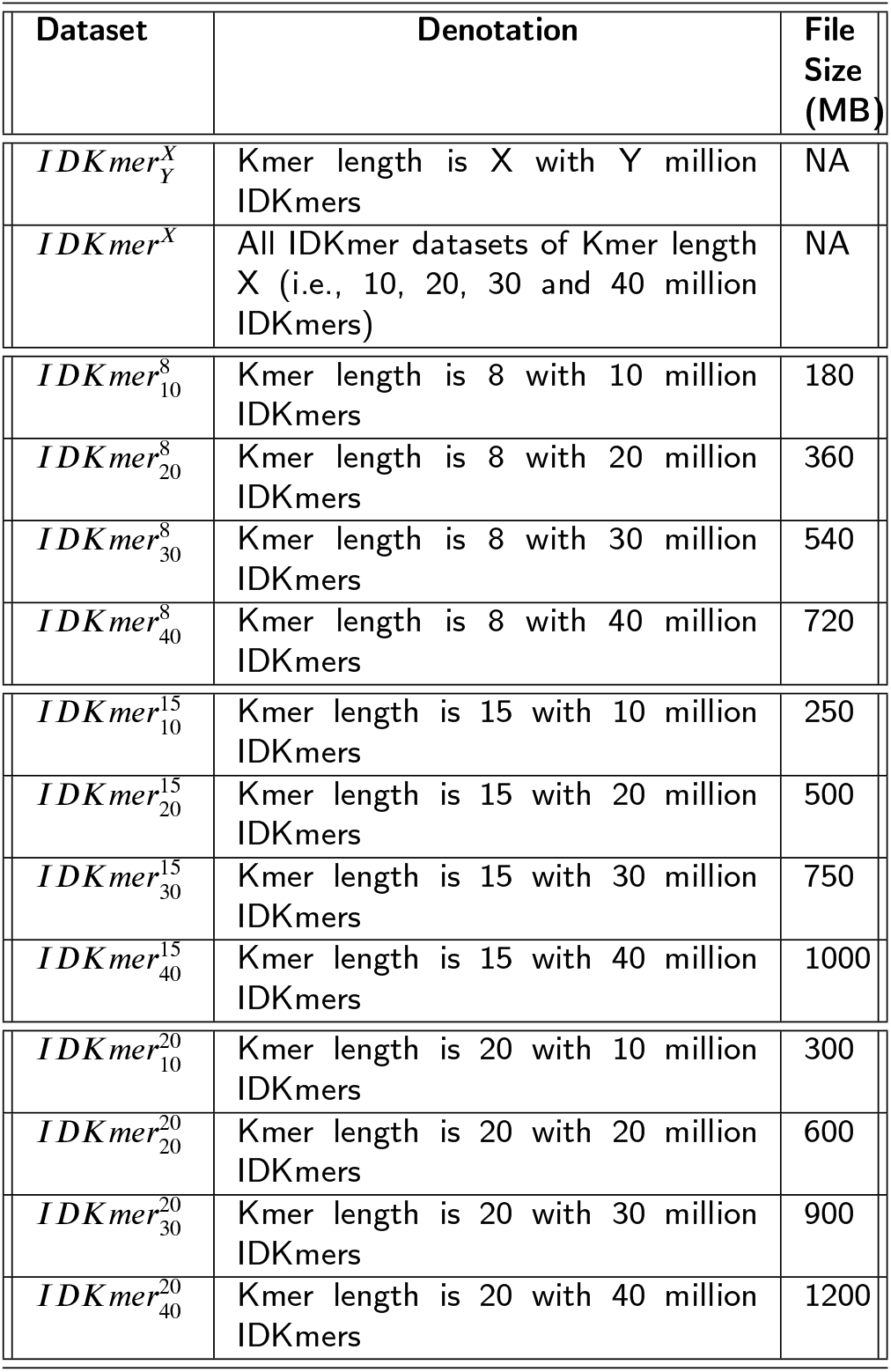
Notation and File size of *IDKmer* Dataset. Mixed Set and Disjoint Set have the same file size. File Size is in MB. NA: Not Applicable.

#### 4.2.2. Biological Network Dataset

Three biological network datasets are considered for real-world data for experimentation: Drug-Gene [dataset] [45], Gene-Disease (Downloaded from [dataset] [46]), and Gene-Gene (Downloaded from [dataset] [46]). In the Drug-Gene dataset, one biological vertex is chemical ID, and another vertex is gene ID. The information provided by the dataset is the Drug-Gene interaction network, i.e., the interaction between genes (proteins encoded by genes) and small molecules. In the Gene-Disease dataset, one biological vertex is gene ID, and the other vertex is disease ID. The dataset provides information regarding the association of disease with genes. This is known as the disorder–gene association [47]. In the Gene-Gene dataset, both the biological vertices are gene IDs. The interaction between gene and gene is called Epistasis [48].

A Mixed Set and Disjoint Set is generated for each biological network dataset. In Disjoint Set, the first vertex is a unique string, and the other vertex is the Gene vertex read from the Drug-Gene dataset. The string of the first edge is “aaaaaaa”, which is incremented alphabetically to have a unique string in each line. Similarly, a unique string is the first vertex of the Disjoint Sets of Gene-Disease and Gene-Gene dataset. The other vertex is the Disease vertex, and Gene vertex values are read from Gene-Disease and Gene-Gene datasets, respectively.

### 4.3. Performance Parameters

Let ℱ𝒫𝒫, 𝒯, 𝒩, ℳℬ, ℬ, and *n* be the false positive probability, time taken (in seconds), number of operations (in million), file size in megabyte (MB), Bloom Filter’s size (in bits), and number of input sequences, respectively. Equation (2), (3), (4), (5) and (6) define accuracy, million operations per seconds (MOPS), megabytes per second (MBPS), second per operation (SPO), and bits per item (BPS), respectively. These parameters are used to determine the performance of BionetBF.

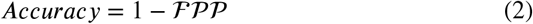

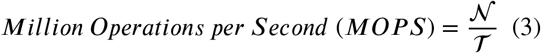

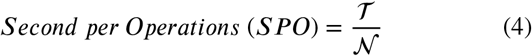

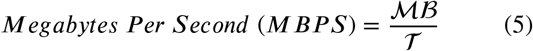

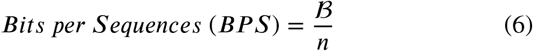

## 5. RESULTS

We have conducted extensive series of tests to validate the performance of BionetBF using diverse datasets: 12 synthetic datasets (Section 4.2 provides the detailed procedure for generation of the dataset) (Table 1 provides the notation and file size details) and biological network dataset (Downloaded from [46]) (Table 2 provides the details of file size and the number of lines). We conducted a rigorous experiment to prove two concerning points of biological networks:

**Table 2.**
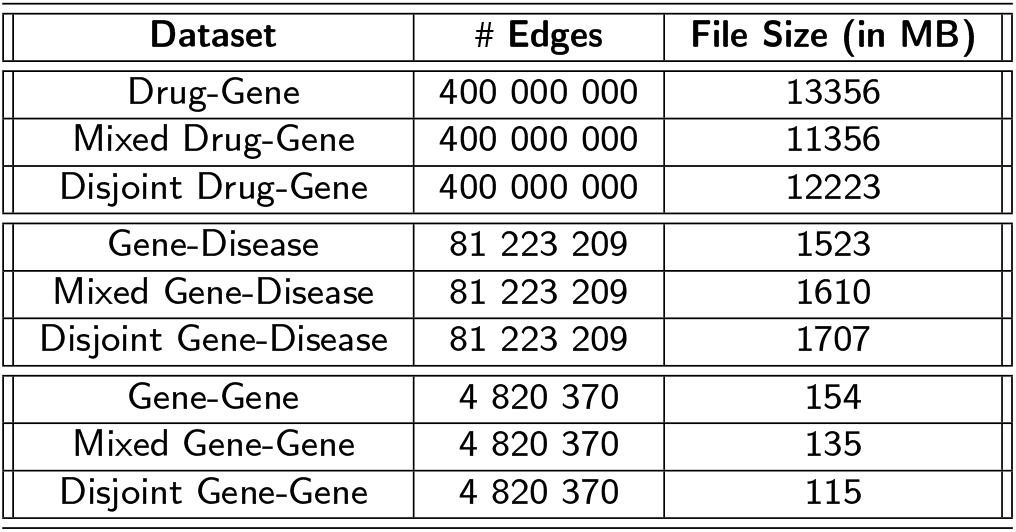
Number of edges and File size of biological network Dataset. # Edges: Number of Edges.

1. BionetBF performs fast processing of the overwhelming sized dataset using tiny amount of memory
2. It has lowest error

BionetBF drastically reduces memory requirements, and we present its analysis and results in this section. We have conducted the experiments in low-cost Ubuntu-Desktop computer with 4GB RAM and Core-i7 processor.

### 5.1. BionetBF

This section presents the analysis of BionetBF with *k* = 2 using the IDKmer dataset. Figure 3a represents the insertion time of BionetBF with different IDKmer datasets. BionetBF takes 1.43 sec for *IDK*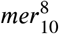 and 7.42 sec for *IDK*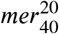 On average, with every increment of 10 million IDKmer insertions, the time increases by 1.41 sec, 1.6 sec, and 1.86 sec for *IDKmer*^8^, *IDKmer*^15^, and *IDKmer*^20^, respectively. Due to the different sequence lengths, the three average values vary, although they have the same number of IDKmers. The IDKmer with a long sequence length requires more reading time. Figure 3b elucidates the time taken by BionetBF after querying the Same Set, Mixed Set and Disjoint Set of different IDKmer datasets. BionetBF for querying Same Set takes 1.47 sec for *IDK* 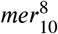 and 7.5 sec for *IDK* 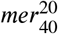. In the Same Set, with every increment of 10 million IDKmer queries, the time increases by 1.46 sec, 1.64 sec, and 1.88 sec for *IDKmer*^8^, *IDKmer*^15^, and *IDKmer*^20^ dataset on average, respectively. BionetBF for querying the Mixed Set takes 1.46 sec for *IDK* 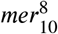 and 7.56 sec for *IDK* 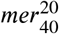 In the Mixed Set, with every increment of 10 million IDKmer queries, the time increases by 1.44 sec, 1.65 sec, and 1.89 sec on average for *IDKmer*^8^, *IDKmer*^15^, and *IDKmer*^20^, respectively. BionetBF for querying Disjoint Set takes 1.46 sec for *IDK* 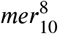 and 7.6 sec for *IDK* 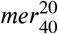. In the Disjoint Set, with every increment of 10 million IDKmer queries, the time increases by 1.43 sec, 1.65 sec, and 1.91 sec on average for *IDKmer*^8^, *IDKmer*^15^, and *IDKmer*^20^, respectively. The insertion or query time of BionetBF is the combination of both reading and execution time. Thus, if the whole dataset is kept in RAM, the operation time can be further increased; however, it is not possible for the biological network dataset because they are usually of massive size. Figure 4 highlights the FPP of the BionetBF. The Same Set gives the number of true positives equal to the number of IDKmers inserted. It indicates the FPP of BionetBF is zero. The *IDKmer*^8^ and *IDKmer*^15^ give zero FPP for both Mixed Set and Disjoint Set. The *IDK* 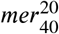 gives FPP of 0.00086 and 0.0017 for Mixed Set and Disjoint Set, respectively.

**Figure 3:**
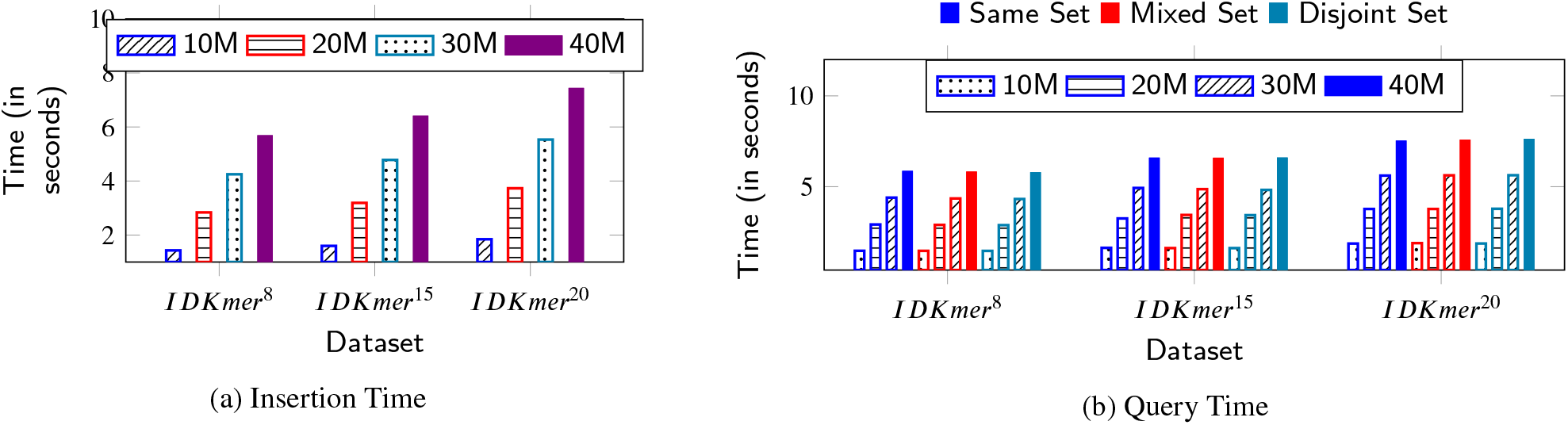
Insertion and Query Time (BionetBF): Comparison of time taken by BionetBF for (a) Insertion Operation and (b) Querying Operation. The Query operation is performed on Same Set, Mixed Set and Disjoint Set. Lower is better. 10M: 10 million, 20M: 20 million, 30M: million, and 40M: 40 million number of IDKmer queries.

**Figure 4:**
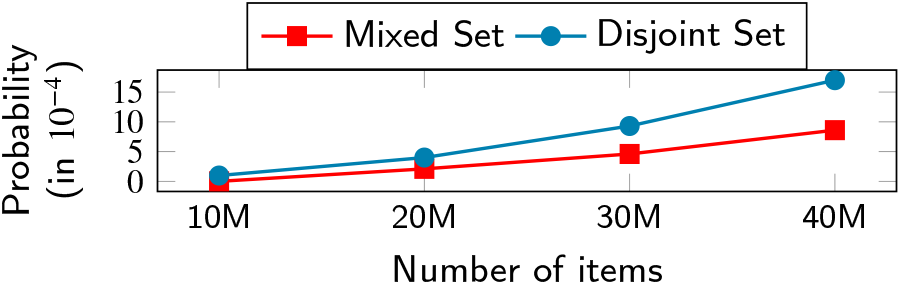
False Positive Probability (BionetBF): Comparison of false positive probability of BionetBF; querying Mixed Set and Disjoint Set. The dataset is IDKmer having sequence length 20. The false positive probability of Mixed Set and Disjoint Set of IDKmer having sequence length 8 and 15 is zero. Lower is better. 10M: 10 million, 20M: 20 million, 30M: million, and 40M: 40 million number of IDKmer queries.

Figure 5 illuminates the accuracy of BionetBF, which is 100% in the case of *IDKmer* ^8^ and *IDKmer*^15^ for both Mixed Set and Disjoint Set. Moreover, the accuracy of the Same Set is 100%. Figure 5a and Figure 5b illustrate the accuracy of the Mixed Set and Disjoint Set, respectively. The accuracy of the Mixed Set and Disjoint Set for *IDKmer*^20^ is 99.91% and 99.83%, respectively. Thus, BionetBF has more than 99% accuracy for a large data file.

**Figure 5:**
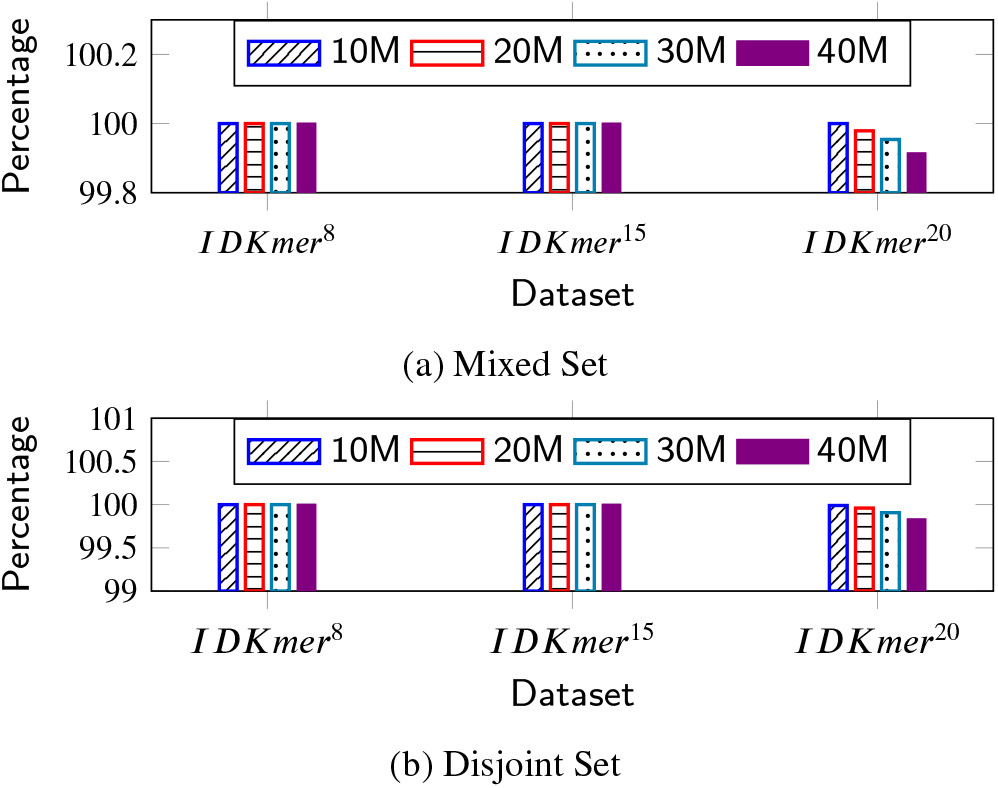
Accuracy (BionetBF): Comparison of accuracy of Same Set, Mixed Set and Disjoint Set queried to BionetBF. Accuracy of Same Set is 100%. Lower value is better. 10M: 10 million, 20M: 20 million, 30M: million, and 40M: 40 million number of IDKmer queries.

Figure 6 represents the performance of the insertion operation. Figure 6a exhibits MOPS where the highest MOPS is 7.06 *(IDK* 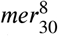 and the lowest is 5.36 *(IDK* 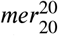 in the IDKmer dataset. Figure 6b evince SPO in which the highest SPO is 1.87 *(IDK* 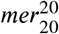 and the lowest is 1.42 *(IDK* 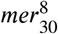). Figure 6c depicts MBPS where the highest MBPS is 162.78 *(IDK* 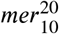 and the lowest is 125.87 *(IDK* 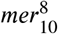). Similarly, Figure 7 delineates the performance of the query operation. Figure 7a, Figure 7b, and Figure 7c exhibit query MOPS for Same Set, Mixed Set and Disjoint Set, respectively. The highest MOPS for Same Set, Mixed Set and Disjoint Set is 6.86 *(IDK* 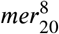), 6.9 *(IDK* 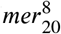), and 6.95 *(IDK* 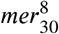), respectively, whereas the lowest MOPS is 5.31 *(IDK* 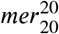), 5.29 *(IDK* 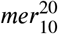), and 5.26 *(IDK* 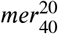), respectively. Likewise, Figure 7d, Figure 7e, and Figure 7f manifest query SOP for Same Set, Mixed Set and Disjoint Set, respectively. The highest SPO for Same Set, Mixed Set and Disjoint Set is 1.89*E* − 07 *(IDK* 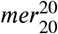), 1.89*E* − 07 *(IDK* 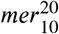), and 1.9*E* − 07 *(IDK* 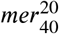), respectively, while the lowest SPO is 1.46*E*−07 *(IDK* 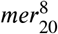), 1.45*E* − 07 *(IDK* 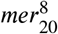), and 1.44*E* − 07 *(IDK* 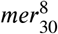), respectively. Furthermore, Figure 7g, Figure 7h, and Figure 7i show the query MBPS for Same Set, Mixed Set and Disjoint Set, respectively. The highest MBPS for Same Set, Mixed Set and Disjoint Set is 160.86 *(IDK* 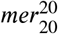), 160.06 *(IDK* 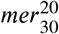), and 160.6 *(IDK* 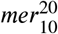), respectively, while the lowest MBPS is 122.7 *(IDK* 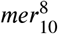), 122.95 *(IDK* 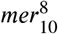, and 123.2 *(IDK* 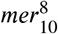), respectively. Another parameter is used to determine the performance of BionetBF, i.e., the number of bits per sequence (BPS) (Equation (6)) illustrated in Table 3. The bits are calculated by the BionetBF data structure size. The BPS is based on two fixed parameters: BionetBF Bloom Filter size and the number of input sequences. Hence, BPS is the same for all IDKmer dataset with the same number of IDKmers. The highest BPS is 10.85 exhibited by the 10 million IDKmers, while the lowest is 2.71 exhibited by the 40 million IDKmers.

**Table 3.** Bits Per Sequences (BPS) for BionetBF

| Number of IDKmers | BPS |
| --- | --- |
| 10 million | 10.85 |
| 20 million | 5.425 |
| 30 million | 3.612 |
| 40 million | 2.71 |

**Figure 6:**
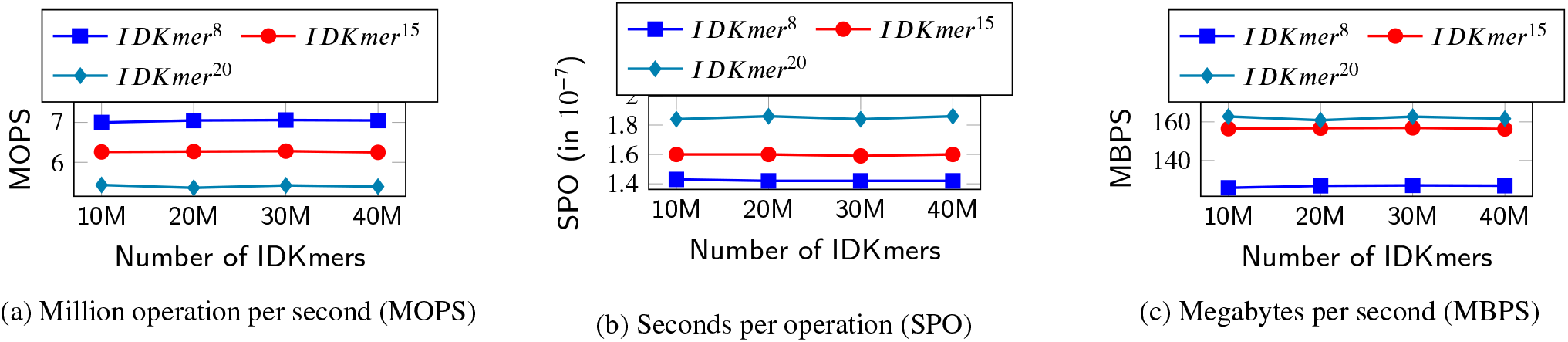
Performance of Insertion Operation (BionetBF): Comparison of performance of BionetBF based on the (a) number of million insertion operations executed per second (MOPS), (b) time taken for execution of one insertion operation (SPO), and (c) execution of Megabyte per second (MBPS) in case of insertion operation. In case of (a) and (c), higher is better. In case of (b), lower is better. 10M: 10 million, 20M: 20 million, 30M: million, and 40M: 40 million number of IDKmers inserted.

**Figure 7:**
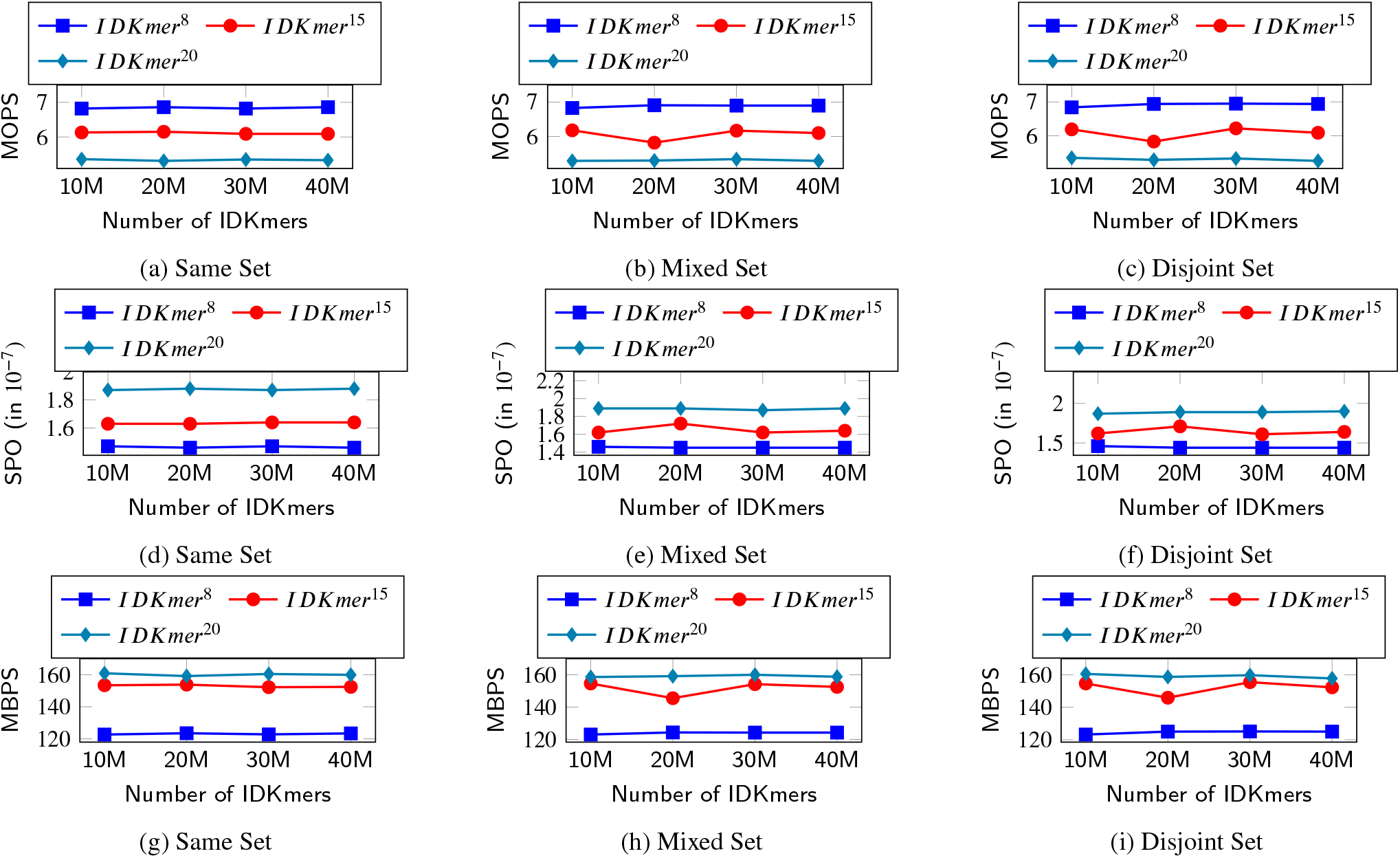
Performance of Query Operation (BionetBF): Comparison of performance of BionetBF based on [(a), (b) and (c)]: the execution of million query operations per sec (MOPS), [(d), (e) and (f)]: the time taken in seconds per operation (SPO), and [(g),(h) and (i)]: the Megabyte executed per second in case of query operation. The query operation is executed on Same Set, Mixed Set and Disjoint Set. In case of [(a), (b) and (c)] and [(g),(h) and (i)], higher is better. In case of [(d), (e) and (f)], lower is better. 10M: 10 million, 20M: 20 million, 30M: million, and 40M: 40 million number of IDKmer queries.

### 5.2. Comparison with Other filters

In this section, BionetBF is compared with Cuckoo Filter [29] (Code available at [49]) and Libbloom (Code available at [50]). The filter size of the Cuckoo Filter is based on the total number of input sequences. In the experiments, the highest number of sequences inserted is 40 million; based on this, the filter size of the Cuckoo Filter is 100MB. If the number of total sequences is reduced, Cuckoo Filter does not insert all sequences in case of *IDKmer*^20^. Cuckoo Filter takes numbers as input; hence, a hash function is added to convert the sequences to numbers. Hence, the total number of hash functions used by the Cuckoo Filter is 3 if no kicking occurs. Another important point is if Cuckoo Filter is executed with the same dataset multiple times, it gives different false positives. We have considered the least number of false positives after executing the Cuckoo Filter multiple times for a single dataset. The Libbloom is the standard Bloom Filter [8]. Similar to Cuckoo Filter, in the case of Libbloom, the total number of sequences considered is 40 million. The memory size is 71 MB, and to achieve an FPP of 0.001, the Libbloom takes 10 hash functions. In the experiment, the filter size of BionetBF is 15MB. Similar to BionetBF, the Kmer ID and sequence of the IDKmer dataset are concatenated and inserted as a single sequence into Cuckoo Filter and Libbloom. Table 4 highlights the filter size and the number of hash functions used by Cuckoo Filter, Libbloom, and BionetBF in the experiment.

**Table 4.** Details of Cuckoo Filter, Libbloom, and BionetBF. #Hash functions: Number of hash functions executed per operation.

| Filter | Filter Size | #Hash functions |
| --- | --- | --- |
| Cuckoo Filter | 100MB | 3 (if no kicking) |
| Libbloom | 71MB | 10 |
| BionetBF | 15MB | 2 |

Figure 8 and Figure 9 show the comparison among Cuckoo Filter, Libbloom and BionetBF based on the time taken for insertion and query operation, respectively. Figure 8 and Figure 9 represent the 10, 20, 30 and 40 million IDKmers, respectively. The Libbloom exhibits the highest time due to the highest number of hash functions as illuminated by Figure 8 and Figure 9. In the case of insertion of *IDK*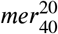, Cuckoo Filter, Libbloom and BionetBF take 12.97 sec, 23.95 sec, and 7.42 sec, respectively. Hence, Libbloom takes the longest time, and BionetBF takes the least time. With every increment of 10 million IDKmers, the insertion time of the Cuckoo Filter increases by on average 2.93, 3.18, and 3.24 sec for *IDKmer*^8^, *IDKmer*^15^, and ^20^, respectively. In Libbloom, the insertion time increases by on average 5.59, 5.88 and 5.99 sec for *IDKmer*^8^, *IDKmer*^15^, and *IDKmer*^20^, respectively. BionetBF takes 1.41, 1.6 sec and 1.86 more for *IDKmer*^8^, *IDKmer*^15^, and *IDKmer*^20^, respectively with increase million IDKmers. Figure 9a, Figure 9b, and Figure 9c elucidate the differentiation of Cuckoo Filter, Libbloom and BionetBF based on the query time of Same Set, Mixed Set, and Disjoint Set, respectively. In Same Set (Figure 9a), the query time taken for *IDK* 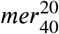 by Cuckoo Filter, Libbloom and BionetBF is 12.4 sec, 20.84 sec, 7.5 sec, respectively, whereas for *IDK*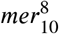 is 2.79 sec, 4.81 sec, and 1.47 sec, respectively. Likewise, in Mixed Set (Figure 9b), the query time taken for *IDK*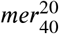 by Cuckoo Filter, Libbloom and BionetBF is 12.38 sec, 17.82 sec, 7.56 sec, respectively; while for *IDK* 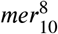 is 2.79 sec, 3.87 sec, and 1.46 sec, respectively. In Disjoint Set (Figure 9c), the query time taken for *IDK* 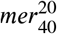 by Cuckoo Filter, Libbloom and BionetBF is 12.47 sec, 14.78 sec, 7.6 sec, respectively, and for *IDK* 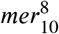 is 2.82 sec, 2.91 sec, and 7.6 sec, respectively. Similar to insertion time, in the case of query time, Libbloom takes the longest time, and BionetBF takes the least time. With every increment of 10 million IDKmer queries, the query time of Cuckoo Filter increases by on average 2.79, 3.04, and 3.12 sec for Disjoint Set of *IDKmer*^8^, *IDKmer*^15^, and *IDKmer*^20^, respectively. In the case of Libbloom, the query time increases by on average 3.31, 3.6 and 3.7 sec for the Disjoint Set of *IDKmer*^8^, *IDKmer*^15^, and *IDKmer*^20^, respectively. BionetBF takes 1.43, 1.65 and 1.91 sec more for the Disjoint Set of *IDKmer*^8^, *IDKmer*^15^, and *IDKmer*^20^, respectively, with every increment of 10 million IDKmer queries.

**Figure 8:**
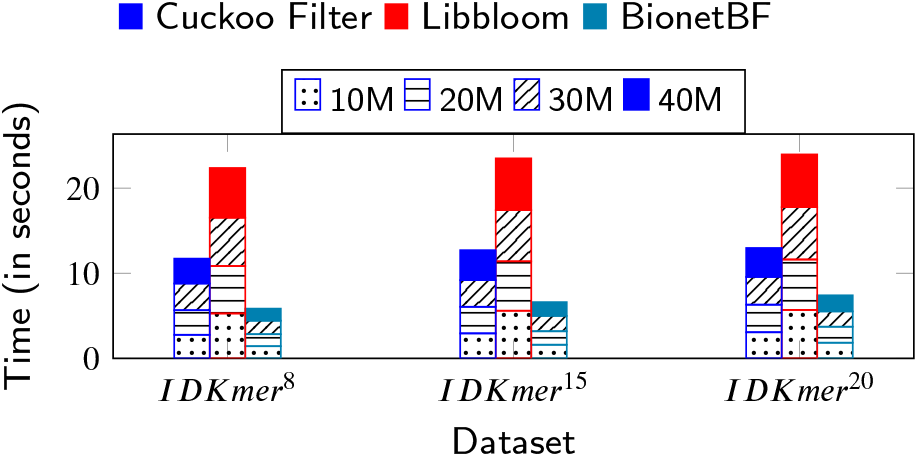
Insertion Time (Cuckoo Filter vs Libbloom vs BionetBF): Comparison of insertion time among Cuckoo Filter, Libbloom, and BionetBF. 10M: 10 million, 20M: 20 million, 30M: 30 million, 40M: 40 million number of IDKmers inserted.

**Figure 9:**
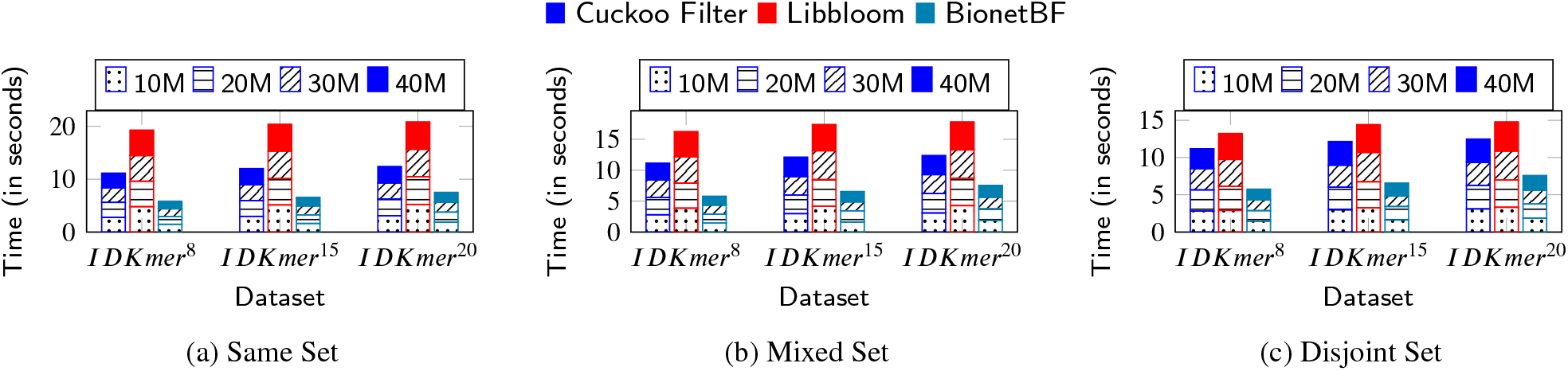
Query Time (Cuckoo Filter vs Libbloom vs BionetBF): Comparison of time taken by Cuckoo Filter, Libbloom and BionetBF for the Same Set, Mixed Set and Disjoint Set for query operation. 10M: 10 million, 20M: 20 million, 30: million, and 40: 40 million number of IDKmers queried.

Figure 10 and Figure 11 delineate the comparison among Cuckoo Filter, Libbloom and BionetBF based on the FPP for Mixed Set and Disjoint Set. Clearly, the Cuckoo Filter has the highest FPP, which is more than the desired FPP, i.e., 0.001 in all IDKmer datasets. Libbloom exhibits less than desired FPP in all IDKmer datasets. In the Mixed Set (Figure 10), Cuckoo Filter has 0.003 (approx.) on average more FPP than BionetBF, and Cuckoo Filter has 0.0029 (approx.) on average more FPP than Libbloom. In the case of Libbloom, BionetBF has 0.0000144 (approx.) on average more FPP. It is an extremely low difference, but considering the size of the filters, BionetBF has better performance. BionetBF has zero FPP in the majority IDKmer datasets, whereas, except for the 10 million IDKmer datasets, Libbloom has some FPP. In the Disjoint Set (Figure11), the Cuckoo Filter has 0.0059 (approx.) on average more FPP than BionetBF, and the Cuckoo Filter has 0.006 (approx.) on average more FPP than Libbloom. In the case of Libbloom, it has 0.0000128 (approx.) on average less FPP than BionetBF. In the case of the Disjoint Set, BionetBF, with its small filter size, outperforms Libbloom. BionetBF has zero FPP in *IDKmer*^8^ and *IDKmer*^15^. Cuckoo Filter has zero FPP in all IDKmer datasets, and Libbloom has zero FPP only in datasets having 10 million IDKmers. Therefore, BionetBF exhibits superior performance in regard to insertion time, query time and FPP with a small filter size for a high number of IDKmers.

**Figure 10:**
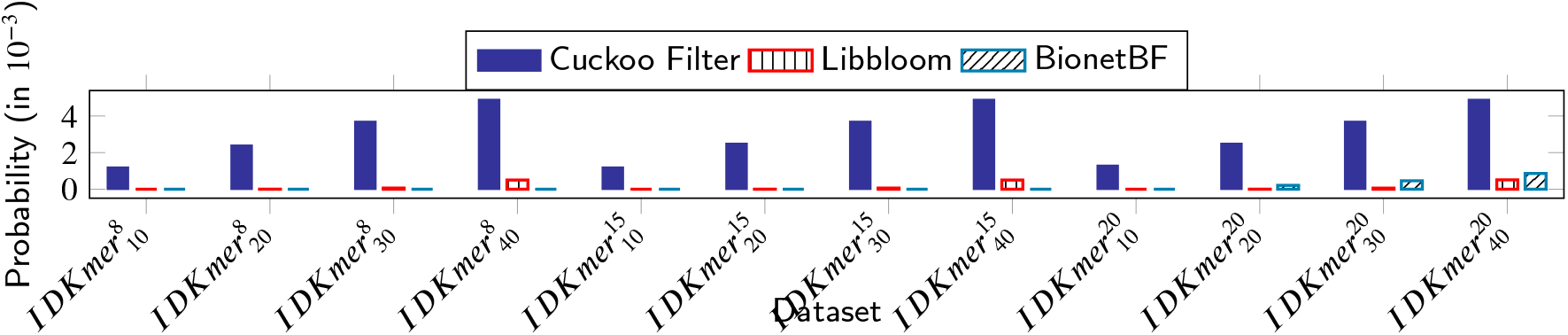
False Positive Probability (Libbloom vs Cuckoo Filter vs BionetBF): Comparison of false positive probability of querying Mixed Set to Libbloom, Cuckoo Filter and BionetBF.

**Figure 11:**
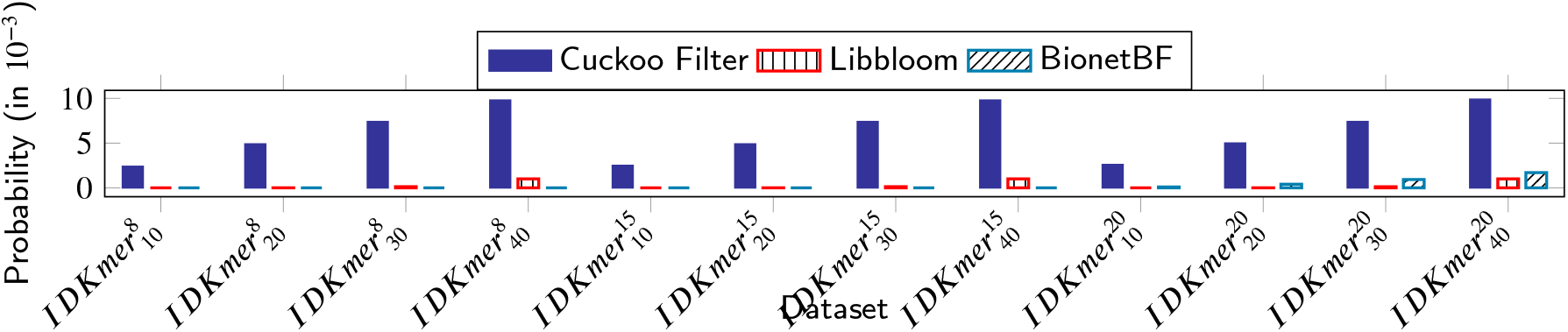
False Positive Probability (Libbloom vs Cuckoo Filter vs BionetBF): Comparison of false positive probability of querying Disjoint Set to Libbloom, Cuckoo Filter and BionetBF.

Figure 12 highlights the analogy among Cuckoo Filter, Libbloom and BionetBF based on the accuracy. In all IDKmer datasets, Libbloom and BionetBF have 100% accuracy, whereas Cuckoo Filter does not have 100% accuracy, but it is still more than 99%. This result is obvious because the data structure of Cuckoo Filter and Libbloom are constructed to achieve the desired FPP.

**Figure 12:**
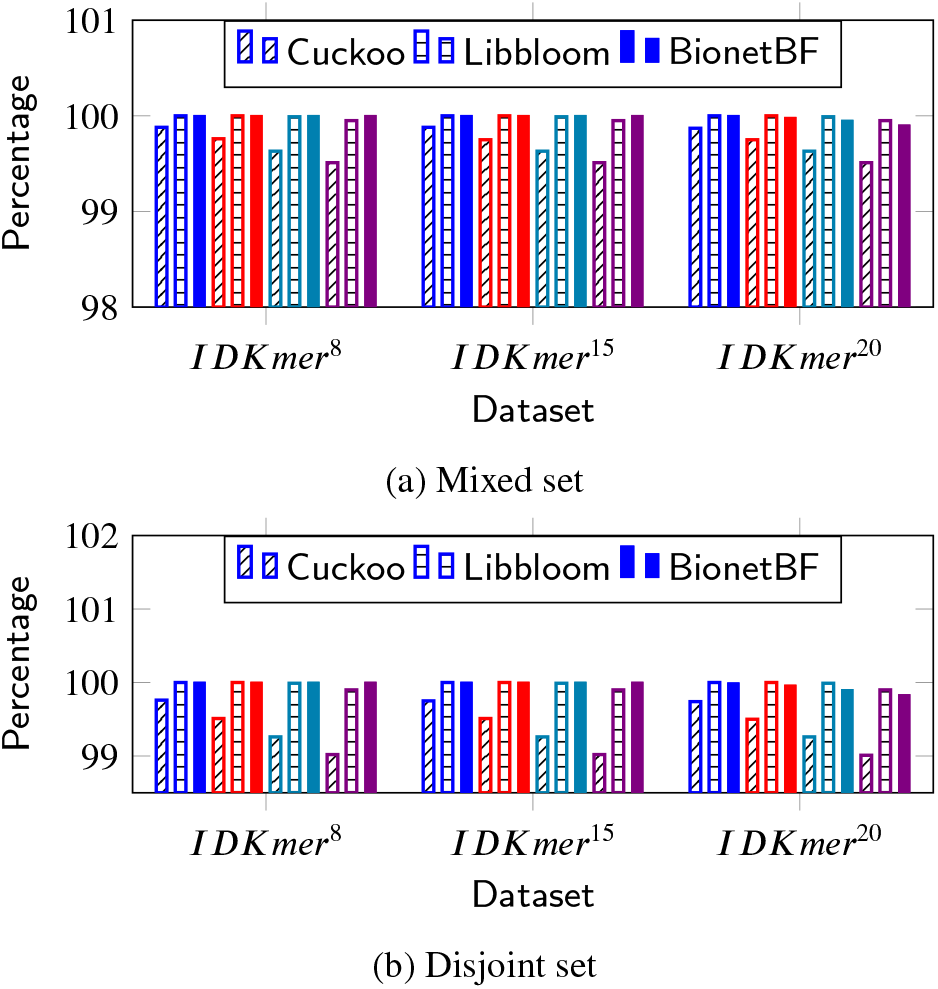
Accuracy (Cuckoo vs Libbloom vs BionetBF): Comparison of accuracy of querying Same, Mixed and Disjoint set to Libbloom, Cuckoo, and BionetBF. Higher value is better. 10M: 10 million, 20M: 20 million, 30M: million, and 40M: 40 million number of IDKmers queried.

Figure 13 delineate comparison among Cuckoo Filter, Libbloom and BionetBF based on the performance of insertion operation. Figure 13a demonstrates the insertion MOPS of Cuckoo Filter, Libbloom and BionetBF. The highest insertion MOPS of Cuckoo Filter, Libbloom and BionetBF is 3.62 *(IDK* 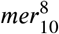), 18.94 *(IDK* 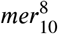), and 7.06 *(IDK* 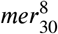), respectively, whereas the lowest MOPS is 3.09 *(IDK* 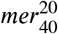), 1.67 *(IDK* 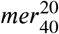), and 5.36 *(IDK* 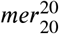), respectively. BionetBF has the highest insertion MOPS with 1.88× and 3.53× (on average) better than Cuckoo Filter and Libbloom; the Cuckoo Filter is 1.88× (on average) better than Libbloom. Figure 13b illustrates the insertion SPO of Cuckoo Filter, Libbloom and BionetBF. The highest SPO of Cuckoo Filter, Libbloom, and BionetBF is 3.24*E* − 07 *(IDK* 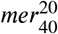), 5.99*E* − 07 *(IDK* 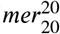), and 1.87*E* − 07 *(IDK* 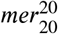) respectively, while the lowest SPO is 2.77*E* − 07 *(IDK* 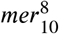), 5.28*E* − 07 *(IDK* 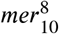), and 1.42*E* − 07 *(IDK* 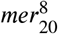), respectively. In insertion SPO, Libbloom is highest, indicating bad performance compared to the other two; BionetBF is approximately 0.54× and 0.29× (on average) higher than Cuckoo Filter, and Libbloom and Cuckoo Filter is approximately 0.53 times (on average) better than Libbloom. Figure 13c highlights the insertion MBPS of Cuckoo Filter, Libbloom and BionetBF. The highest MBPS of Cuckoo Filter, Libbloom and BionetBF is 97.66 *(IDK* 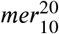), 52.77 *(IDK* 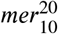), and 162.78 *(IDK* 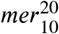), respectively, whereas the lowest MBPS is 61.36 *(IDK* 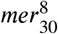), 32.19 *(IDK* 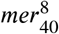), and 125.87 *(IDK* 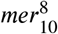), respectively. In the case of MBPS, BionetBF is highest with approximately 1.88× and 3.53× (on average) better than Cuckoo Filter and Libbloom; the Cuckoo Filter is approximately 1.88× (on average) better than Libbloom. Similar to the performance of insertion operation, BionetBF has better performance in the query operation. Figure 14, Figure 15, and Figure 16 highlight the analogy among Cuckoo Filter, Libbloom and BionetBF based on the performance of query operation. Figure 14 represent the query MOPS of Cuckoo Filter, Libbloom and BionetBF where Figure 14a, Figure 14b, and Figure 14c highlights the query MOPS for Same Set, Mixed Set and Disjoint Set, respectively. In Same Set (Figure 14a), the highest MOPS shown by Cuckoo Filter, Libbloom and BionetBF is 3.59 *(IDK* 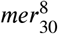), 2.08 *(IDK* 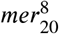), and 6.86 *(IDK* 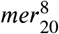), respectively, whereas the lowest MOPS is 3.2 *(IDK* 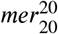), 1.9 *(IDK* 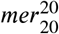), and 5.31 *(IDK* 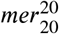), respectively. In Mixed Set (Figure 14b), the highest MOPS is Cuckoo Filter, Libbloom and BionetBF is 3.59 *(IDK* 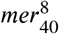), 2.58 *(IDK* 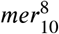), and 6.9 *(IDK* 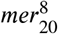), respectively, while the lowest MOPS is 3.2 *(IDK* 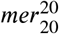), 2.25 *(IDK* 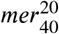), and 5.29 *(IDK* 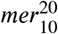), respectively. In Disjoint Set (Figure 14c), the highest MOPS is Cuckoo Filter, Libbloom, and BionetBF is 3.58 *(IDK* 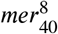), 3.44 *(IDK* 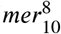), and 6.95 *(IDK* 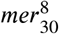), respectively, and the lowest MOPS is 3.19 *(IDK* 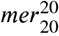), 2.71 *(IDK* 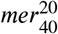), and 5.26 *(IDK* 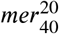), respectively. In MOPS, BionetBF is approximately 1.8×, 1.8× and 1.81× (on average) better than Cuckoo Filter in the Same Set, Mixed Set and Disjoint Set respectively, and approximately 3.07×, 2.56× and 2.048× (on average) better than Libbloom in Same Set, Mixed Set and Disjoint Set, respectively. The Cuckoo Filter is approximately 1.704×, 1.42×, and 1.13× (on average) better than Libbloom in Same Set, Mixed Set and Disjoint Set, respectively. Figure 15 shows the comparison of performance based on SPO for query operation. Figure 15a, Figure 15b and Figure 15c illustrate the query SPO for Same Set, Mixed Set and Disjoint Set, respectively. In Same Set (Figure 15a), the highest SPO in Cuckoo Filter, Libbloom and BionetBF is 3.12*E* − 07 *(IDK* 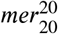), 5.23*E* − 07 *(IDK* 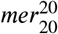), and 1.89*E* − 07 *(IDK* 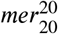), respectively, whereas the lowest SPO is 2.79*E* − 07 *(IDK* 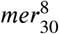), 4.81*E* − 07 *(IDK* 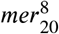), and 1.46*E* − 07 *(IDK* 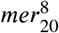), respectively. In Mixed Set (Figure 15b), the highest SPO in Cuckoo Filter, Libbloom and BionetBF is 3.12*E* − 07 *(IDK* 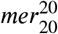), 4.45*E* – 07 *(IDK* 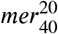), and 1.89*E* − 07 *(IDK* 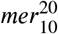), respectively, whereas the lowest SPO is 2.79*E* − 07 (*IDK* 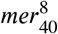), 3.87*E* − 07 *(IDK* 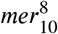), and 1.45*E* − 07 *(IDK* 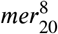), respectively. In Disjoint Set (Figure 15c), the highest SPO in Cuckoo Filter, Libbloom and BionetBF is 3.13*E* − 07 (*IDK* 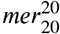), 3.7*E* − 07 *(IDK* 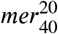), and 1.9*E* − 07 *(IDK* 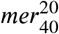), respectively, whereas the lowest SPO is 2.79*E* − 07 *(IDK* 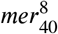), 2.91*E* − 07 *(IDK* 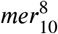), and 1.44*E* − 07 *(IDK* 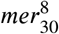), respectively. In SPO, BionetBF is approximately 0.5×, 0.56× and 0.55× (on average) better than Cuckoo Filter in the Same Set, Mixed Set and Disjoint Set, respectively, and approximately 0.33×, 0.39×, 0.49× (on average) better than Libbloom in Same Set, Mixed Set and Disjoint Set, respectively. The Cuckoo Filter is 0.59×, 0.7×, and 0.89× (on average) better than Libbloom in the Same Set, Mixed Set and Disjoint Set, respectively. Figure 16 demonstrates the differentiation among Cuckoo Filter, Libbloom, and BionetBF using performance based on query MBPS. Figure 16a, Figure 16b and Figure 16c illuminate the query MBPS for Same Set, Mixed Set and Disjoint Set, respectively. Figure 16a represents the query MBPS of Same Set where the highest MBPS in Cuckoo Filter, Libbloom, and BionetBF is 96.96 *(IDK* 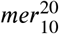), 57.56 *(IDK* 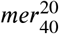), and 160.86 *(IDK* 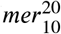), whereas the lowest MBPS is 63.99 *(IDK* 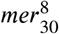), 37.32 *(IDK* 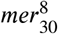), and 122.7 *(IDK* 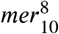), respectively. Figure 16b depicts the query MBPS of Mixed Set where the highest MBPS in Cuckoo Filter, Libbloom, and BionetBF is 96.96 *(IDK* 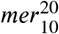), 70.13 *(IDK* 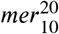), and 160.06 *(IDK* 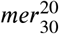), while the lowest MBPS is 64 *(IDK* 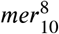), 44.3 *(IDK* 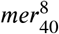), and 122.95 *(IDK* 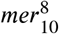), respectively. Figure 16c demonstrates the query MBPS of Disjoint Set where the highest MBPS in Cuckoo Filter, Libbloom, and BionetBF is 96.59 *(IDK* 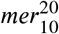 90.2 *(IDK* 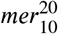), and 160.6 *(IDK* 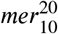), whereas the lowest MBPS is 63.46 *(IDK* 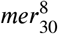), 54.41 *(IDK* 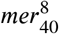), and 123.2 *(IDK* 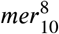), respectively. In MBPS, BionetBF is approximately 1.8×, 1.8× and 1.81× (on average) better than Cuckoo Filter in the Same Set, Mixed Set and Disjoint Set, respectively, and approximately 3.07×, 2.56× and 2.048× (on average) better than Libbloom in Same Set, Mixed Set and Disjoint Set, respectively. The Cuckoo Filter is approximately 1.704×, 1.42×, and 1.13× (on average) better than Libbloom in Same Set, Mixed Set and Disjoint Set, respectively. Moreover, Table 5 highlights the comparison of performance among Cuckoo Filter, Libbloom, and BionetBF based on BPS. Cuckoo Filter occupies approximately 7.42× and 1.4× more BPS compared to Libbloom and BionetBF, respectively. Libbloom occupies approximately 5.3× more BPS than BionetBF.

**Table 5.**
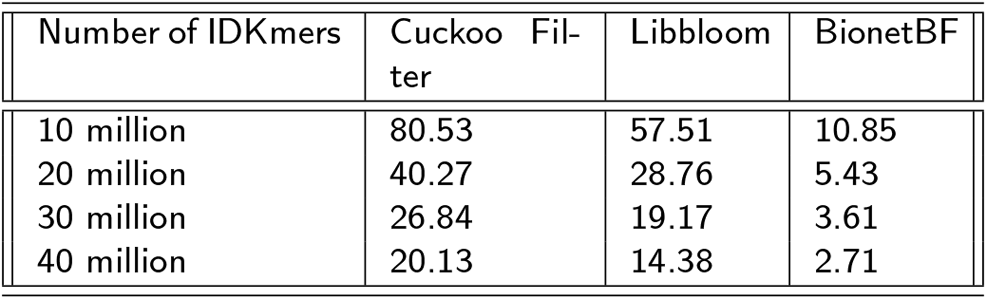
Comparison of memory usage among Cuckoo Filter, Libbloom, and BionetBF based on Bits Per Sequences (BPS).

**Figure 13:**
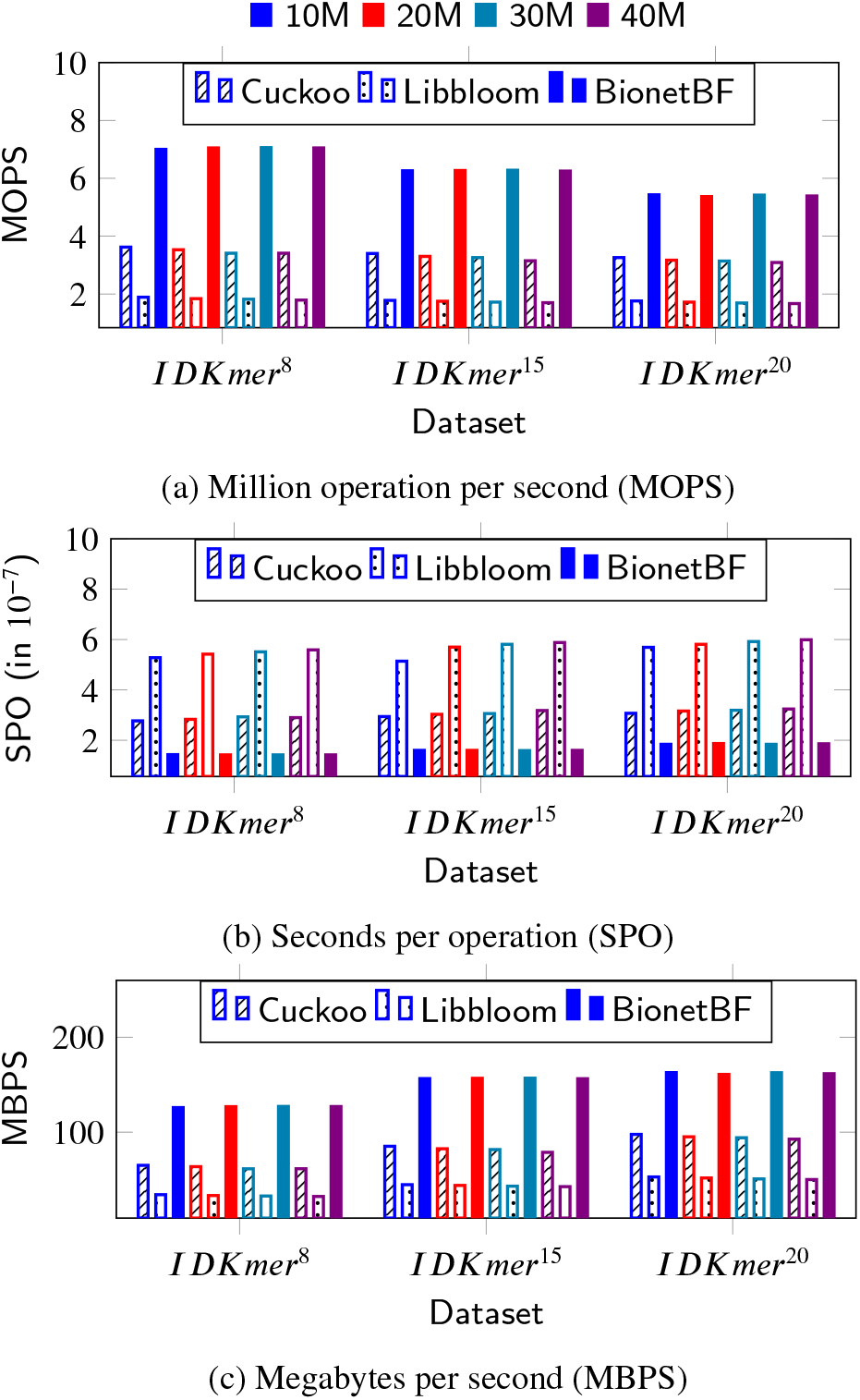
Performance of Insertion Operation (Cuckoo vs Libbloom vs BionetBF): Comparison of performance among Cuckoo, Libbloom, and BionetBF based on the (a) number of million insertion operations executed per second (MOPS), time taken for execution of one insertion operation (SPO), and (c) execution of Megabyte per second (MBPS) in case of insertion operation. In case of (a) and (c), higher is better. In case of (b), lower is better. 10M: 10 million, 20M: 20 million, 30M: million, and 40M: 40 million number of IDKmers queried.

**Figure 14:**
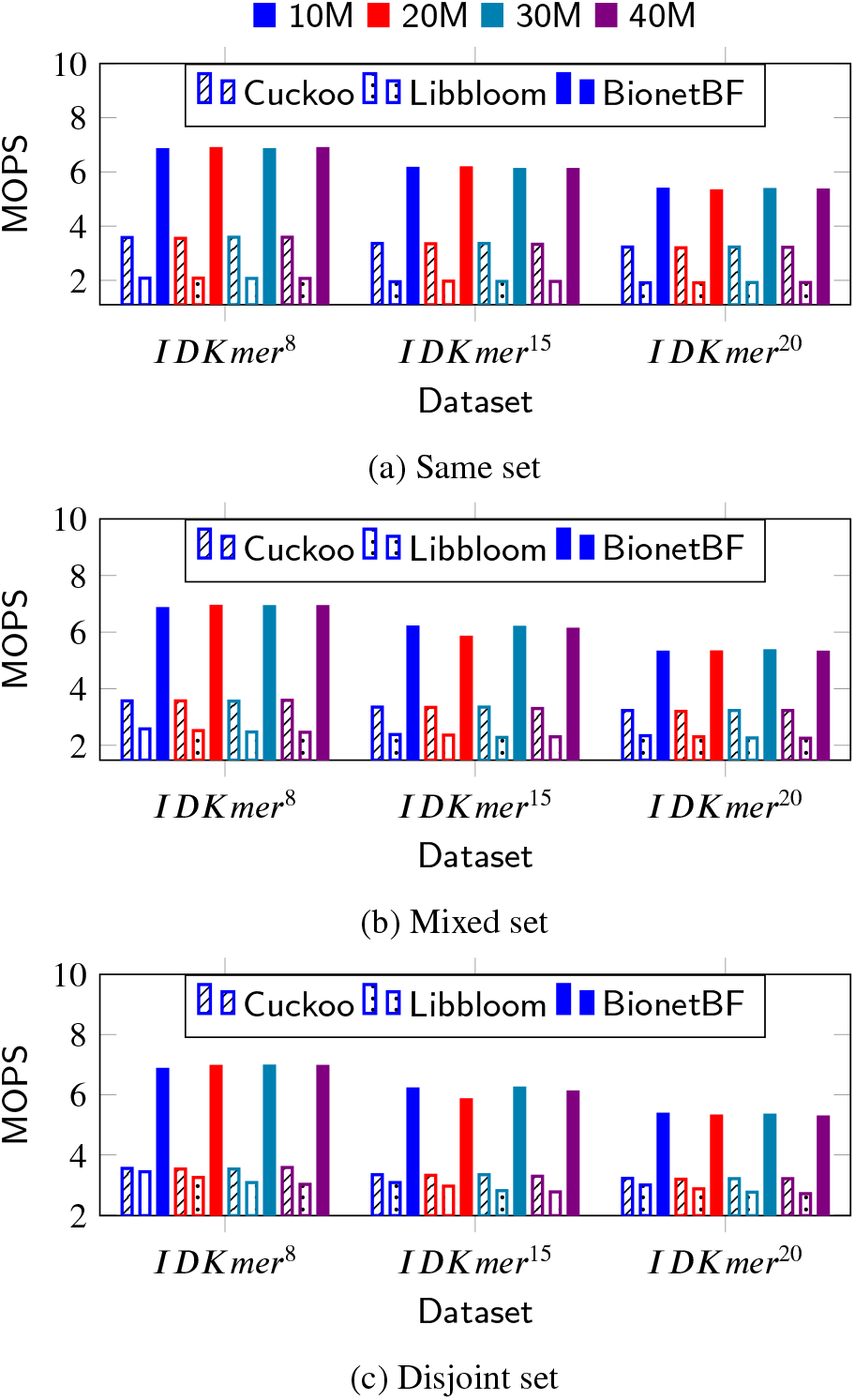
Query MOPS (Cuckoo vs Libbloom vs BionetBF): Comparison of performance among Cuckoo, Libbloom, and BionetBF based on the million query operations executed per second (MOPS) for (a) Same set, (b) Mixed set, and (c) Disjoint set. Higher is better. 10M: 10 million, 20M: 20 million, 30M: million, and 40M: 40 million number of IDKmers queried.

**Figure 15:**
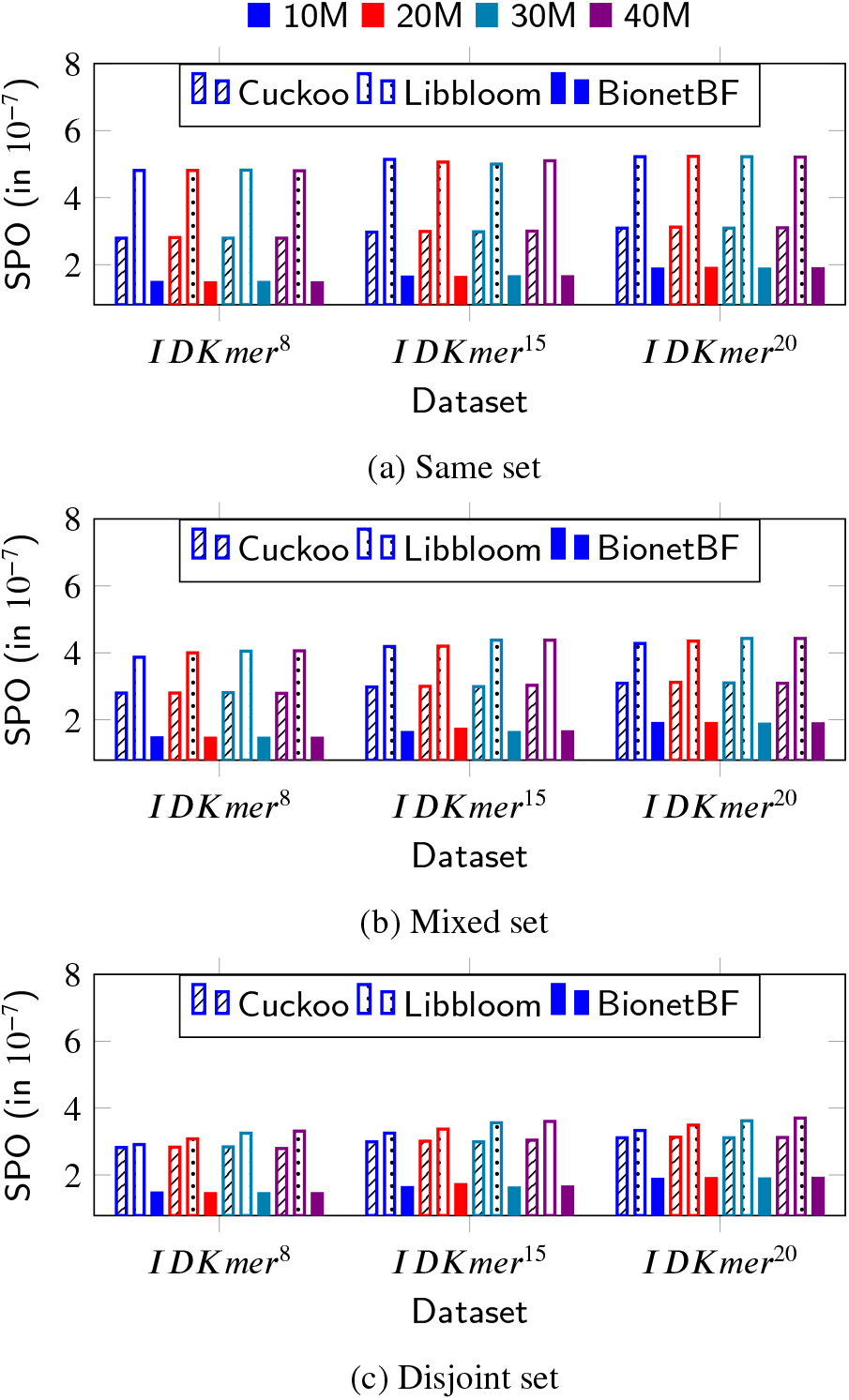
Query SPO (Cuckoo vs Libbloom vs BionetBF): Comparison of performance among Cuckoo, Libbloom, and BionetBF based on second per operation (SPO) for (a) Same set, (b) Mixed set, and (c) Disjoint set. Lower is better. 10M: 10 million, 20M: 20 million, 30M: million, and 40M: 40 million number of IDKmers queried.

**Figure 16:**
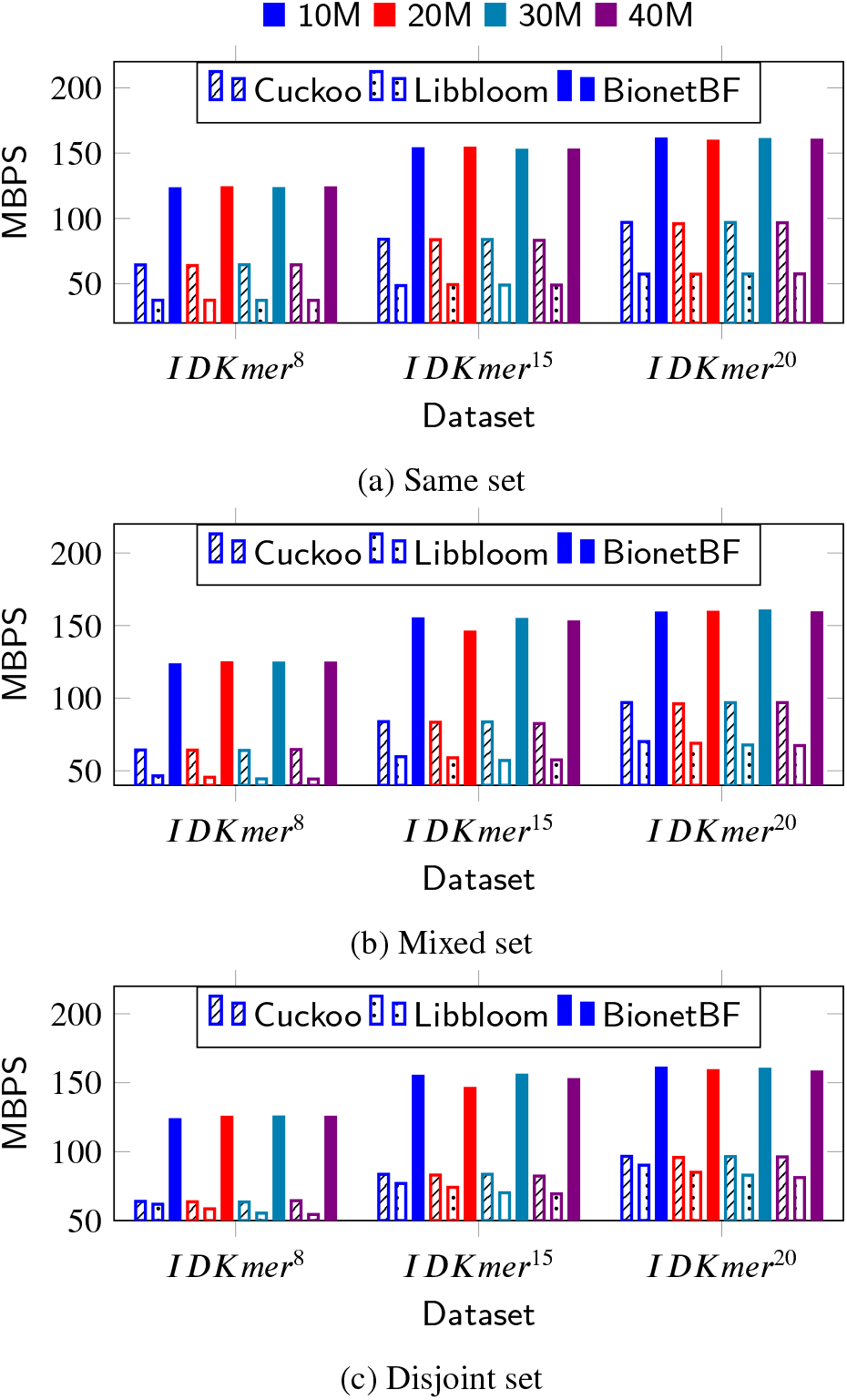
Query MBPS (Cuckoo vs Libbloom vs BionetBF): Comparison of performance among Cuckoo Filter, Libbloom, and BionetBF based on Megabyte per second for query operations in case of (a) Same set, (b) Mixed set, and (c) Disjoint set. Higher is better. 10M: 10 million, 20M: 20 million, 30M:million, and 40M: 40 million number of IDKmers queried.

From these experiments, we have proved the high efficiency and performance of BionetBF compared to other filters, namely, Cuckoo Filter and Libbloom. Another important point is that a small-sized BionetBF performs better than a big-sized Cuckoo Filter and Libbloom. Another filter is also considered for comparison with BionetBF, namely, Xor filter [42] (Code available at [51]). For faster processing, the Xor filter saves all the sequences in an array which is given as input for the construction of the filter. This reduces the time duration of the whole operation, i.e., insertion and query. Usually, genomic data is huge; inserting the whole genomic data in a single array is not possible. Moreover, the Xor filter has a fixed upper limit on the number of repetitive data possible in the dataset. The genomic data are highly repetitive in nature. This condition of the Xor filter is a huge constraint for the genomic data. Therefore, the XOR filter is not appropriate for genomic data.

### 5.3. BionetBF with Biological network

This section provides an analysis and result of experimentation performed on BionetBF using three biological network datasets: Drug-Gene, (b) Gene-Disease, and (c) Gene-Gene. Some details regarding the dataset are mentioned in Section 4 4.2.2. While experimenting with the Drug-Gene dataset, the biological edge (i.e., chemical and gene) are concatenated and inserted or queried to BionetBF. Similarly, in the Gene-Disease dataset, the biological edge (i.e., gene and disease) are concatenated. In the case of Gene-Gene, the interacting genes are concatenated to insert or query to BionetBF. An important point to highlight is the size of the Drug-Gene dataset is 13.4 GB (approx.), Gene-Disease is 1.5GB (approx.), and Gene-Gene is 154MB.

Figure 17 represents the insertion and query time of BionetBF with the biological network dataset. Drug-Gene, Gene-Disease, and Gene-Gene take 76.6 sec, 11.64 sec and 0.88 sec, respectively, to insert biological edges. Similarly, for Disjoint Set, the query time taken by Drug-Gene, Gene-Disease, and Gene-Gene is 76.75 sec, 11.93 sec, and 0.74 sec, respectively. Figure 18 illustrates the FPP of BionetBF with biological network datasets: Drug-Gene and Gene-Gene. The Gene-Gene dataset has zero FPP. Both Drug-Gene and Gene-Disease datasets are big-sized datasets; however, the FPP is very low. Figure 19 illuminates the accuracy of BionetBF with biological network datasets: Drug-Gene (Figure 19a), Gene-Disease (Figure 19b) and Gene-Gene (Figure 19c). The Gene-Gene dataset has 100% accuracy; Drug-Gene and Gene-Disease have an accuracy of more than 99.9%. Figure 20 elucidates the performance of BionetBF with the biological network datasets for both insertion and query operations. Considering both insertion and query operations, BionetBF with the biological network datasets exhibit more than 5 MOPS, below 2.10^−7^ SPO, and more than 130 MBPS. Table 6 illustrates the bits per sequence of the three biological network datasets.

**Table 6.** BPS of biological network

| Dataset | BPS |
| --- | --- |
| Drug-Gene | 0.27 |
| Gene-Disease | 1.34 |
| Gene-Gene | 22.51 |

**Table 7.**
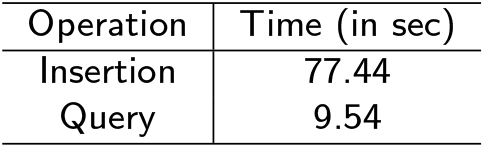
Case Study: Time taken during insertion of Drug-Gene dataset and query of Chemical-gene dataset. Zero number of interactions found.

**Figure 17:**
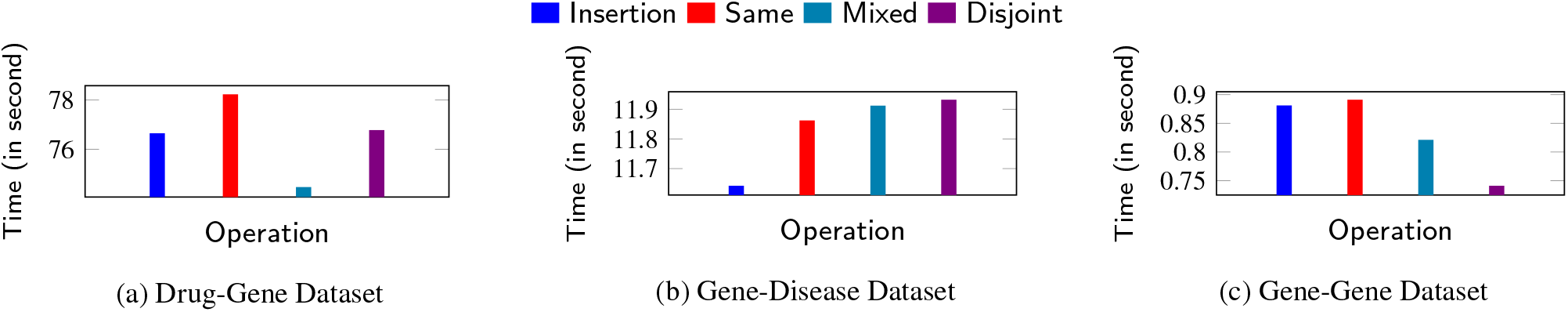
Insertion and Query Time (Biological network Dataset): Time taken for Insertion and Query operation by BionetBF with biological network Dataset:- (a) Drug-Gene, (b) Gene-Disease, and (c) Gene-Gene. Lower is better.

**Figure 18:**
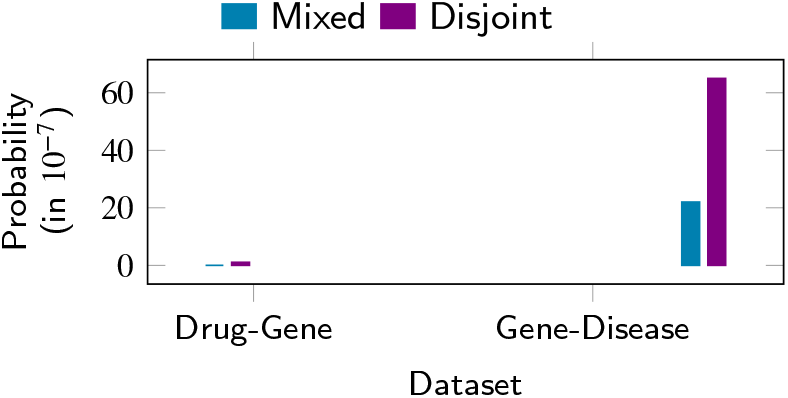
False Positive Probability (Biological network Dataset): False positive probability of BionetBF with biological network dataset: (a) Drug-Gene and (b) Gene-Disease. Lower is better. Gene-Gene dataset has zero false positive probability.

**Figure 19:**
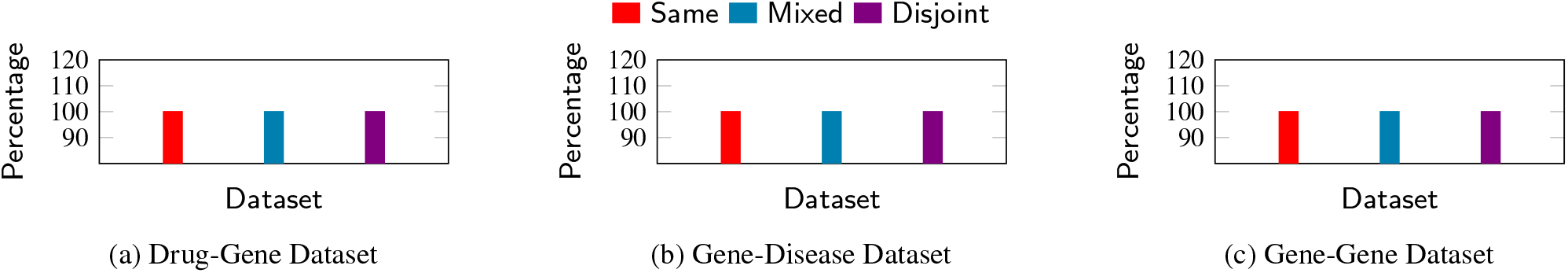
Accuracy (Biological network Dataset): Accuracy of BionetBF with biological network dataset:- (a) Drug-Gene, (b) Gene-Disease, and (c) Gene-Gene. Higher is better.

**Figure 20:**
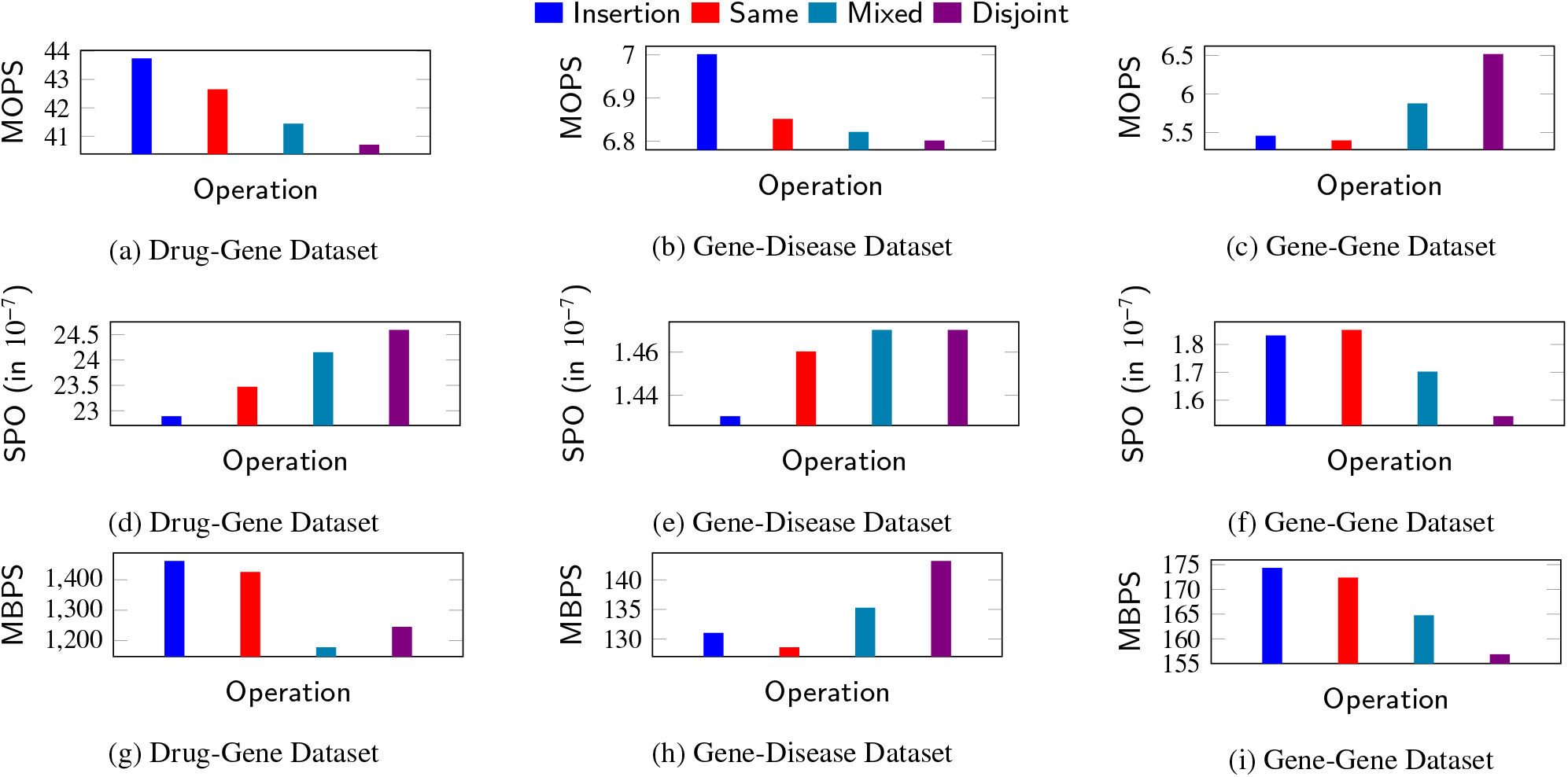
Performance (Biological network dataset): Performance of BionetBF with biological network dataset: [(a), (d), and (g)] Drug-Gene, [(b), (e), and (h)] Gene-Disease, and [(c), (f), and (i)] Gene-Gene, based on [(a), (b), and (c)]: Million operation per second (MOPS). [(d), (e), and (f)]: Seconds per operation (SPO). [(g), (h), and (i)]: Megabyte executed per second (MBPS). In case of [(a), (b) and (c)] and [(g),(h) and (i)], higher is better. In case of [(d), (e) and (f)], lower is better.

### 5.4. Drug-Gene Interaction: A Case Study

Boolean query/membership identification is an important operation in Big Data. Boolean query operation determines whether the queried data is a member of the huge dataset. In the case of Big Graph, boolean query operation determines whether the queried edge is connected to the Big Graph. The huge size of the Big Graph is increasing the complexity of this simple operation which either returns true or false as a response. This section presents a case study to present an application of BionetBF for a faster boolean query. We have considered two real dataset: Drug-Gene dataset and Chemical-Gene dataset (Downloaded from [dataset] [46]).The Drug-Gene and Chemical-Gene datasets have biological data on the interaction between chemicals and genes. To avoid confusion, one dataset is called Drug-Gene and another Chemical-Gene datasets. The size of BionetBF is kept the same as in the above experimentation. The details of the Drug-Gene dataset are mentioned in Section 4.2. The Drug-Gene dataset is 13.4 GB with 400 million drug-gene interactions. The Drug-Gene dataset has a size of 1.14 GB with 62816502 drug-gene interactions. The Chemical-Gene dataset is inserted into the BionetBF, and the Drug-Gene dataset is queried to BionetBF. The insertion time is 77.44 sec, and the query time is 9.54 sec. For every query, BionetBF returned false. In other words, BionetBF was able to respond to around 62.8 million Drug-Gene interactions that are absent in the biological network having 400 million drug-gene interactions within 10 secs.

## 6. Discussion

BionetBF can help in drug discovery. Another implementation is the deduplication of the biological edge. This section discusses some implementation of BionetBF in the biological network field.

One implementation of BionetBF is faster identification of biological edges in a biological network consisting of a particular target protein. The biological networks are stored in the database. Multiple BionetBFs are constructed for each biological network. The size of the BionetBF is based on the size of the biological network. A large biological network with thousands or millions of nodes can be stored in a small-sized BionetBF. It is illustrated in the Result section. Hence, constructing a BionetBF is not much of an overhead. When a target protein and its other interacting protein are determined, the biological edge is checked in BionetBFs in parallel. If a BionetBF returns true, check its related biological network in the database for further processing. Searching the biological network in the database is difficult because searching a Big Graph is an NP-hard problem. However, BionetBF can confirm the membership of a Big Graph within seconds. Another application of BionetBF in drug discovery is security. The biological network can be stored in BionetBF and provided to others without disclosing the main structure of the biological network. Suppose a pharmaceutical company wants to study the compatibility of their drug with another drug belonging to another pharmaceutical company. The latter company does not want to completely provide the information about its drug (say, *DrugA*). However, they also want to encourage the development of the former company’s drug (say *DrugB*). *DrugB* aims to determine whether consumption of *DrugB* along with *DrugA* is safe. *DrugA* company can store the protein-protein interactions in the BionetBF. Also, other information such as chemical-protein interaction can be stored in the same BionetBF. This BionetBF is sent to the *DrugB* company. BionetBF is itself an encrypted data; extracting complete information from BionetBF is difficult. Even using a Brute force attack on the BionetBF, i.e., continuously querying the possible proteinprotein interactions, is very difficult with such a huge number of protein possibilities. *DrugB* can determine its protein-protein interactions and check in the BionetBF whether it is present in Drug A.

Bloom Filters are mostly known for deduplication. BionetBF also an excellent choice for deduplication of repetitive insertion or query of biological edge. While determining the biological network of a biological process, the same biological edge is generated and forwarded to the database for storage. However, the repetition of such biological edge can also be in thousands. Storage of huge repetitive information in databases is a huge issue in Bioinformatics. But, searching and removal of such repetitive biological data is another difficult task. Hence, in such a situation, BionetBF can be implemented. After the generation of the biological edge, check it in the BionetBF. In case it is present, ignore the storage in the database; otherwise, it is stored in the BionetBF first and then forwarded to the database for storage. Another advantage of using BionetBF is slow storage in the database without halting the processing. Databases are in secondary storage; hence, the storage time complexity is more. The application does not have to wait for the database to complete the insertion of the biological data; rather BionetBF can continue working for the membership checking, and the newly biological edge is stored in a buffer and slowly inserted into the database.

## 7. Conclusion

Our proposed system, BionetBF, identifies the membership of biological edge effectively and efficiently. It can store a huge volume of the biological edge using a tiny amount of memory. As we know, biological data are significantly large, requiring huge-sized memory (RAM) to process. Therefore, BionetBF offers an effective and efficient solution for the same. In addition, our proposed solution requires 2.71 bits per biological edge to store in the RAM. Therefore, it proves that it can store mammoth-sized biological data in a few MB of main memory. BionetBF can process the biological data at the pace of 162.78 MBPS in a low-cost computer. Therefore, BionetBF will be able to process much higher MBPS in a higher-end computing system. We conducted a series of experiments using 12 synthetic datasets and three real biological network datasets to validate our proposed method. Experimentally, it is illustrated that the execution time of all the operations is low while mitigating the false positive probability to the lowest. Furthermore, BionetBF has a zero error rate in a few datasets. We compared BionetBF with another two filters: Cuckoo Filter and Libbloom (standard Bloom Filter), where a smaller-sized BionetBF exhibits higher performance than the other two filters. Therefore, BionetBF can enhance the entire system performance of Drug Discovery, Genome coding, etc., due to its low memory footprint, nearly 100% accuracy, and high performance. Notably, BionetBF can dramatically boost any system’s performance. Our experimental results show the processing capabilities of biological data in low-cost hardware. Therefore, it can drastically reduce the computing cost of the entire application system.

BionetBF can be used to store an extensive biological network with thousands or millions of nodes. It can be implemented for quick membership checking of neighbouring nodes of a biological network for a specific protein. Moreover, it features a small memory footprint that can hold information about the huge biological network. Hence, an application can maintain multiple BionetBFs that store nodes of biological networks for different proteins. In drug discovery, BionetBF can be used for passing information securely. BionetBF saves the biological network without disclosing the main structure. BionetBF is an outstanding choice for the deduplication of repetitive biological edges. BionetBF offers constant time complexity, which is ideal for the repetitive insertion or query of biological edge. Thus, it can be a game-changer for genomics, proteomics, and Bioinformatics.

## Acknowledgements

Authors sincerely acknowledge the BigCyber Researh Lab for providing the sufficient resource to carry out the entire research.

## Availability of data and materials

The source code is available at https://github.com/patgiri/BionetBF. The code is written in C programming language. All data are available at the given link. Moreover, we have also added a source code to generate synthesis data from a given data source.

## Footnotes

1 The synthetic dataset size is big, hence cannot be uploaded with the code. However, a sample dataset is provided for example (https://github.com/patgiri/BionetBF). It also includes the synthetic data generation code and the DNA sequence dataset.

